# Cellular stress in brain organoids is limited to a distinct and bioinformatically removable subpopulation

**DOI:** 10.1101/2022.03.11.483643

**Authors:** Ábel Vértesy, Oliver L. Eichmueller, Julia Naas, Maria Novatchkova, Christopher Esk, Meritxell Balmaña, Sabrina Ladstaetter, Christoph Bock, Arndt von Haeseler, Juergen A. Knoblich

## Abstract

Organoids enable disease modeling in complex and structured human tissue, *in vitro*. Like most 3D models, they lack sufficient oxygen supply, leading to cellular stress. These negative effects are particularly prominent in complex models, like brain organoids, where they can prevent proper lineage commitment. Here, we analyze brain organoid and fetal single cell RNA sequencing (scRNAseq) data from published and new datasets totaling over 190,000 cells. We describe a unique stress signature found in all organoid samples, but not in fetal samples. We demonstrate that cell stress is limited to a defined organoid cell population, and present Gruffi, an algorithm that uses granular functional filtering to identify and remove stressed cells from any organoid scRNAseq dataset in an unbiased manner. Our data show that adverse effects of cell stress can be corrected by bioinformatic analysis, improving developmental trajectories and resemblance to fetal data.

## Introduction

Organoids are 3D stem cell cultures that enable human tissue modeling with unprecedented structure and complexity (Eiraku et al., 2008; Kadoshima et al., 2013; Lancaster et al., 2013; Paşca et al., 2015; Qian et al., 2016). At the same time, single cell transcriptomics has become widely used for their characterization. Alongside these recent technological breakthroughs, it has become clear that 3D tissue culture is affected by limited oxygen and nutrient supply to the center of the tissue. As most models lack functional vascularization (Garreta et al., 2021), and therefore rely on limited passive transport across the tissue, diffusion-limited hypoxia is an intrinsic problem in organoids. Nutrient-, and in particular oxygen-limitation are long-known phenomena in tissue models (Malda et al., 2007; Volkmer et al., 2008). Oxygen restriction causes widespread metabolic changes by activating the hypoxia-, glycolysis-, and ER stress-pathways, furthermore it affects differentiation and proliferation (Kültz, 2005; Mohyeldin et al., 2010).

Brain organoids are among the most complex and physically largest organoids and are therefore most affected by the limited nutrient supply of the center (Qian et al., 2019). Nevertheless, this problem has only been recently addressed in brain organoids (Bhaduri et al., 2020; Giandomenico et al., 2019; Mansour et al., 2018; Qian et al., 2020) and its extent is still debated.

It therefore remains an open question if stress is a global or a local issue, thus, how far it affects the 3D tissue culture model. A recent paper claimed that *in vitro* conditions lead to a pervasive stress across the whole organoid, causing immaturity, misspecification, and dissimilarity to fetal tissue (Bhaduri et al., 2020). These observations are in contrast to the previous understanding of spatially limited stress (Qian et al., 2019). This raises the question: how should we handle the data affected by an artificial stress signature? It is unclear if stress is an ‘acute’ signature on top of a cell’s original state, or if stress is leading to a completely different cell fate. While the same stress pathways are also active in the fetal brain, reports disagree on whether it is equivalent to those observed *in vitro* (Bhaduri et al., 2020; Gordon et al., 2021).

Experimental solutions emerged in protocols which increase convection with bioreactors, orbital shakers, or microfluidics. Despite these efforts, 3D cultures above ∼500 μm radius develop a necrotic core with healthy tissue limited to the surface ∼100-300 μm. Further developments involve organoid implantation *in vivo* resulting in subsequent vascularization (Mansour et al., 2018), section culture (Giandomenico et al., 2019; Qian et al., 2020), or bioengineering solutions (Garreta et al., 2021). These approaches aim to increase nutrient supply, but neither is currently as scalable as the standard organoid culture which therefore remains the mainstay of organoid research. Until a widely applicable and scalable experimental solution emerges, tissue health and cellular stress persist as a problem for the field.

While most large single-cell RNA-seq studies on diverse brain organoid systems reported glycolytic or ER stressed clusters (Kanton et al., 2019; Tanaka et al., 2020; Velasco et al., 2019), there is no consensus on how to identify them, what is happening in these cells, and what are the consequences of stressed cells on the organoid? To measure the prevalence and consequences of stress in brain organoids, we analyzed differentiation, maturation, and identity of ∼160,000 single cells from newly presented and published cortical and cerebral organoid (together: brain organoids) datasets.

We found stressed cells in all organoid samples, forming a distinct subpopulation. Stressed cells showed widespread transcriptional changes beyond stress pathway activity, we therefore call it ‘stressed-state’. We do not find this stress-state *in vivo*, therefore it is likely an artifact. Eliminating artificial cell populations is essential to truly recapitulate *in vivo* conditions. As stressed cells are currently unavoidable, we developed *granular functional filtering* (Gruffi), an unbiased computational algorithm to isolate stressed cells. Gruffi can clarify developmental trajectories and increase similarity of *in vitro* fetal datasets.

## Results

### A distinct population of ER stressed- and glycolytic-cells exist in all analyzed organoids

We reanalyzed recent, landmark single cell transcriptomics studies and performed new experiments (Bhaduri et al., 2020; Eichmüller et al., 2022; Kanton et al., 2019; Khan et al., 2020; Velasco et al., 2019) (**Fig 1A**) to answer 3 questions: What stress pathways are active in organoids? Does stress occur in all or only certain organoid protocols? Is cellular stress limited to a group of cells or is it pervasive?

**Fig. 1:**
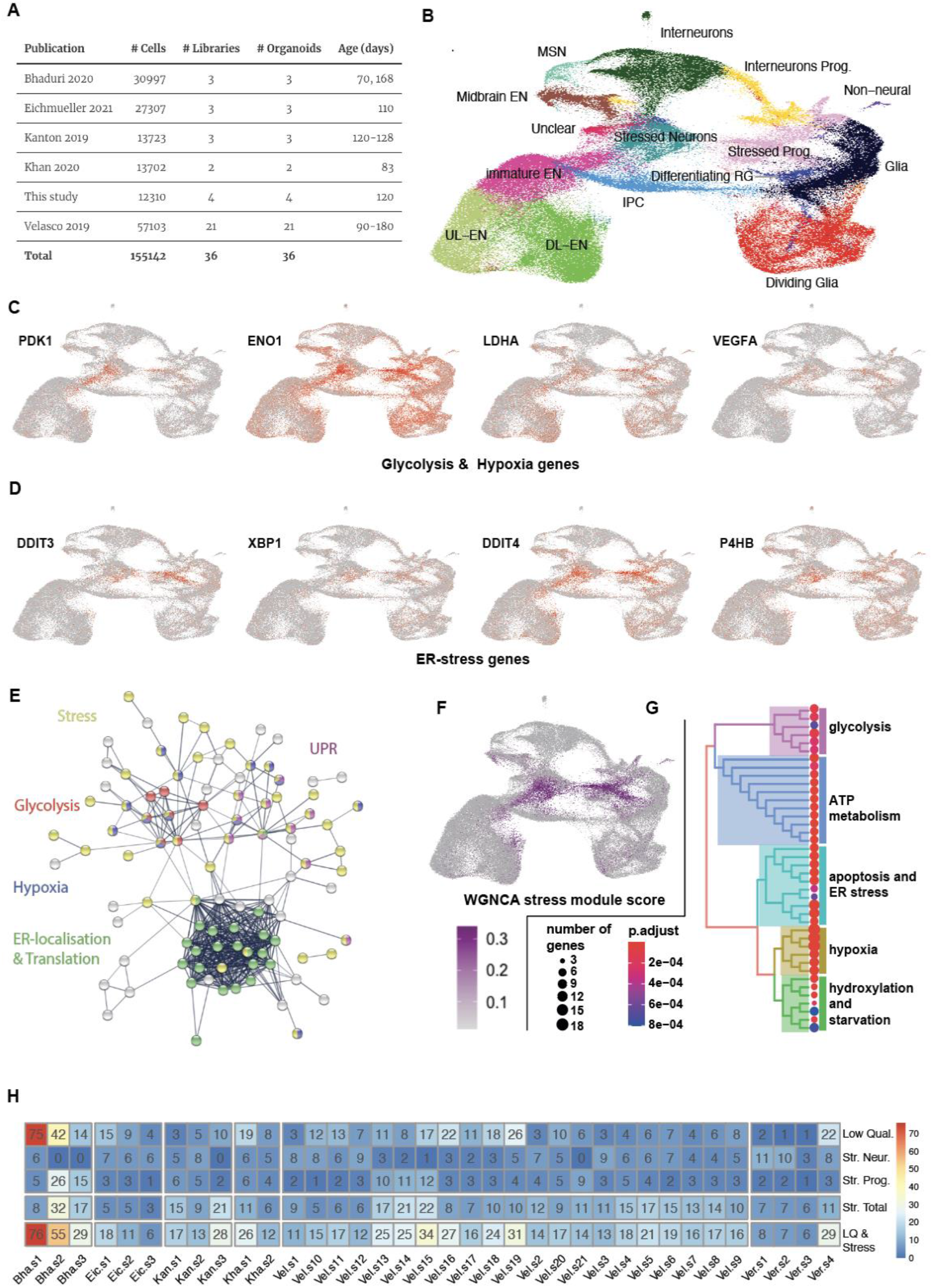
A distinct population of ER-stressed- and glycolytic-cells exist in all analyzed organoids. (**A**) The list of samples and datasets analyzed in this study encompasses mature cortical organoids from multiple key publications. (**B**) UMAP embedding of the integrated dataset. Clustering with cell type annotation shows the expected neural cell types, but also reveals two stressed subpopulations. (**C**) Key marker genes for glycolysis, hypoxia or (**D**) ER-stress are specifically enriched in stress clusters. (**E**) Protein-protein interaction map of GO-term enrichments on the top 150 stressed-cluster enriched genes (by log fold change). Highlighted terms: Cellular response to stress (red, GO:0033554, 2e-07, 0.47); Response to hypoxia (blue, GO:0001666, 3e-10, 0.9); Response to unfolded protein (yellow, GO:0006986, 2e-06, 0.97); Glycolytic process (limegreen, GO:0006096, 1e-05, 1.36); Protein localization to endoplasmic reticulum (cyan, GO:0070972, FDR: 2e-20, strength 1.35) – covering nearly the same ribosomal genes as: Translational initiation (GO:0006413, 2e-20, 1.35). (**F**) WGCNA analysis (see Appendix Figure S1 for other modules) of variable genes identifies a gene module specific to stressed cells, (**G**) which is enriched in stress related terms (**H**) Percentage of low quality cells; cells in stressed-neuron and -progenitor clusters and their union, quantified across all datasets.

We included all those mature, wild type samples that were prepared on the 10X Chromium single - cell platform starting from .fastq files using the same pipeline (methods). To focus on stress in the neural lineage, we removed all cells that are not part of brain development and are a result of mispatterning, sometimes observed in organoids. For a proportional representation of datasets, we sub-sampled ∼160K from a total of 300K cells.

Cellular stress can lead to a perturbation of essential processes, thus affecting cell quality in scRNAseq. Therefore, we applied a minimal filtering, keeping all cells with >500 genes, less than 20% mitochondrial- and 30% ribosomal-reads (Ilicic et al., 2016; Luecken and Theis, 2019). This resulted in a median depth of 3651 UMI/cell (methods). We integrated and analyzed the resulting datasets in Seurat (v4), and found the previously reported cell types (**Fig 1B**). The UMAP separated dividing cells and glia cells from neurons (horizontally) and excitatory-from interneurons (vertically). Besides, there were multiple clusters in the center of the UMAP, which were less well defined by marker gene expression.

### Stress is a common hallmark of the two largest unidentified clusters

Differential gene expression and QC-metric analysis revealed that the ‘unidentified’ central clusters consisted of cells uniquely expressing stress markers and low-quality cells (**Fig 1C, Fig EV1A** and **B**). The stress genes were part of endoplasmic reticulum (ER) stress: *CHOP* (or *DDIT3*), *XBP1, DDIT4, P4HB* (Rashid et al., 2015); glycolysis (*ENO, HK2, PGK1, GAPDH*), and hypoxia: *PDK1, PHD, GLUT1* (or *SLC2A1*) (Lee et al., 2020)(**Fig 1C** and **Fig EV1C-J**).

To better understand the nature of stress in these cells, we analyzed all significantly enriched genes in the stress-clusters. We found that, in either cluster, more than half of the top 50 coding genes were part of ‘response to stress’ (GO:0006950), and apoptosis related terms were among the strongest enriched terms (Table S1). The biggest enriched stress pathways were ‘regulation of cell death’ (GO:0010941), ‘response to hypoxia’ (GO:0001666), and ‘response to endoplasmic reticulum stress’ (GO:0034976). Surprisingly, metabolic terms were both among the strongest and largest enriched terms, highlighting that metabolic shift is a hallmark of stressed cells in organoids.

We then calculated the GO-term enrichment within the 150 strongest enriched coding genes of both stress clusters together, and visualized these on the protein-protein interaction (PPI) map (**Fig 1E**, methods). We highlighted enriched GO-terms (FDR < 5e-7) forming connected PPI clusters, revealing the interplay of glycolysis, hypoxia, unfolded protein response, and translation with the general stress response (**Fig 1E**, Table S1). To identify all genes co-regulated with stress, we applied scWGCNA (single cell weighted gene co-expression network analysis, (Morabito et al., 2021)) and found 12 gene modules (Appendix Figure S1A), one of which was specific to stressed cells (Stress module **Fig 1F**). Gene set enrichment analysis (GSEA) on the stress module identified the strongest enrichment for “response to hypoxia”, “cellular response to ER stress”, and GO-terms of glycolytic processes (**Fig 1G**, Appendix Figure S1B). To test whether stress occurred in all samples, we quantified the contribution of each dataset to the stress clusters. We found that all datasets contained stressed cells, to a generally high (median 13%), but highly variable fraction (50% CV, **Fig 1F**). Thus, stressed cells are a general feature of organoids regardless of conditions, lab of origin or protocol used.

While the initial clustering-based approach identified cells with stress signatures, it had three major limitations. First, while some clusters are too large and comprise mixed populations (Appendix Figure S1C), others can be too small to find marker genes by differential gene expression analysis (DGEA). Second, cluster boundaries are often not well-defined, especially when dealing with developmental trajectories. Third, the resulting limitations in DGEA obstruct the identification of stress genes so that results vary by the dataset and parameters used. Together, these limitations affecting DE could explain why previous studies identified disparate gene sets, like ‘Glycolytic cells’ in (Kanton et al., 2019; Nowakowski et al., 2017) vs. ‘ER stressed cells’ in (Bhaduri et al., 2020; Tanaka et al., 2020). To overcome these issues, we tested different clustering resolutions (Appendix Figure S1D). As none of these could separate the distinct populations of cells within ‘Stressed Neurons’ (**Fig 1B**), we concluded that a new approach is needed to identify and exclude stressed cells.

### Granular functional filtering identifies stressed cells unbiasedly

#### Functional scoring highlights cellular stress regardless of cluster boundaries

To universally identify stressed cells, we established a sample- and data-independent definition for stressed genes. Using gene lists from well-characterized pathways defined as GO-terms (methods), we aggregated information from all genes per pathway by an expression-scoring method widely used for cell cycle scoring (Tirosh et al., 2016). Therein, we downloaded gene lists per GO-term from Ensembl, calculated their average expression and normalized it to randomly sampled control genes of matching expression level (methods). Finally, we evaluated if functional scoring helps to characterize stressed cells. We found that high scores for ‘glycolytic process’ (GO:0006096) and ‘response to endoplasmic reticulum stress’ (GO:0034976), were the strongest signatures of stress-clusters (**Fig 2A**) and provided clearer separation between stressed and non-stressed cells than cluster boundaries. Importantly, high scores marked mostly overlapping cell populations. The coactivation of additional scores, such as ‘response to starvation’ (GO:0042594, **Fig EV1F**) and ‘cellular response to hypoxia’ (GO:0071456, **Fig EV1G**) corroborated a complex stress-identity. Comparing neuronal and glial cell types revealed that all non-dividing glial cells showed higher ER stress scores (**Fig 2B**). We therefore designed our algorithm to accommodate for cell-type specific background when identifying stressed cells.

**Fig. 2:**
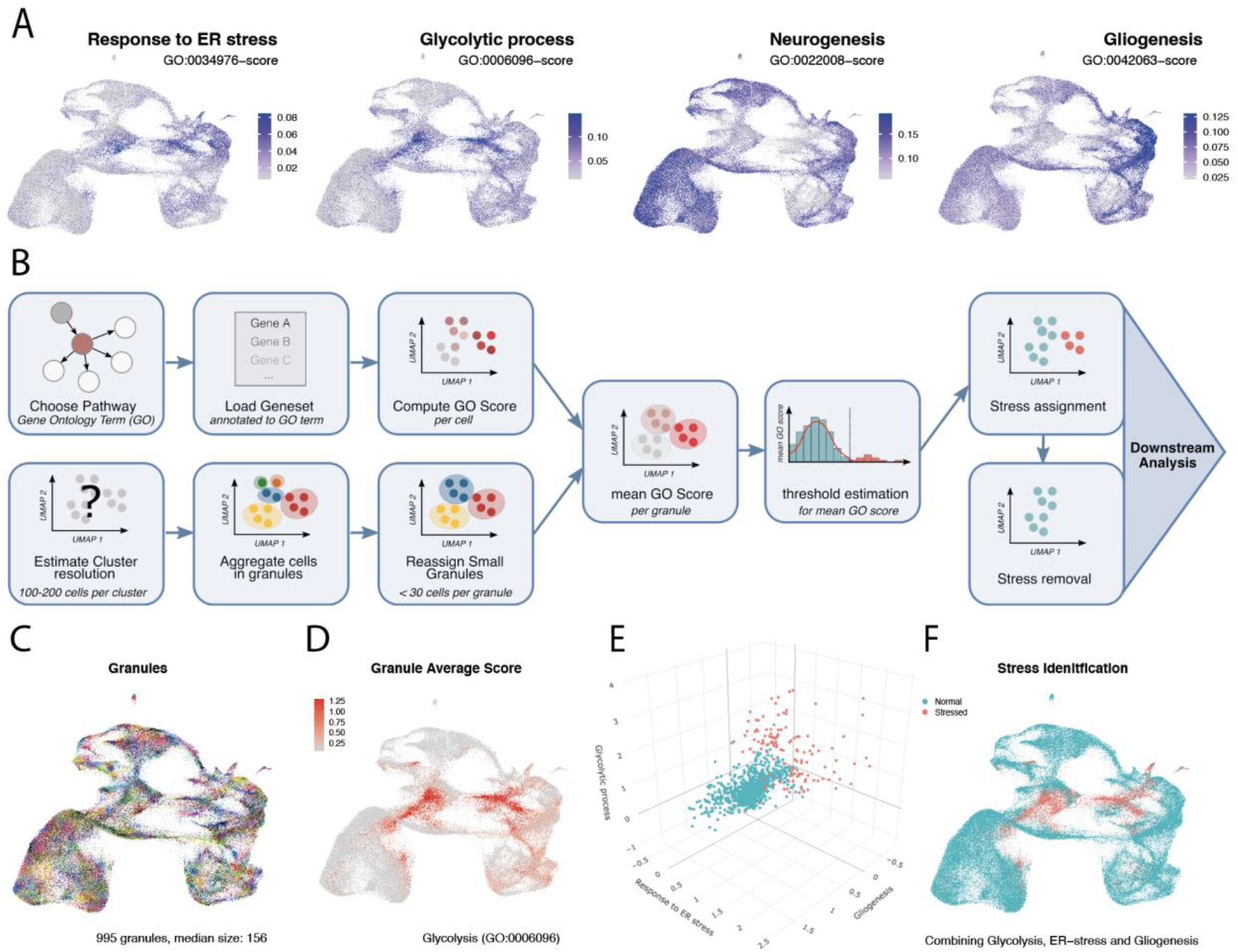
Granular functional filtering identifies stressed cells unbiasedly. Granular functional filtering identifies stressed cells unbiasedly (**A**) Gene-set scores per cell for the two strongest stress signatures (ER-stress and glycolysis) and the two cardinal processes in the developing brain (neurogenesis and gliogenesis). The complementary expression signatures suggest a mutually exclusive neural- or stressed-fate. (**B**) Overview of Gruffi’s stress classification. After preprocessing steps including the computation of PCA and UMAP embeddings, a Gene Ontology pathway is selected, respective gene sets are retrieved, and per cell GO-scores are calculated. At the same time, an ideal clustering resolution is estimated, such that cells are assigned to granules of (in median) ∼100-200 cells, and small clusters (<30) are reclassified. Next, to overcome high variability and detection noise caused by single cell resolution, average and cell number normalized granule scores are calculated, and respective score-thresholds are estimated based on the scores dispersion of the first peak. Finally, stressed granules are identified by a combination of scores, and isolated from the dataset for separate analysis or dataset cleaning and further downstream analysis is possible.(**C**) Gruffi defined 995 granules by snn clustering containing a median of 156 cells. (**D**) Granule granule scores for glycolysis shown on UMAP. (**E**) 3 dimensional stress score threshold estimation by Gruffi using default setting, requiring high glycolytic and ER-stress, low gliogenesis score to define stressed cells. (**F**) Stressed cells classified by Gruffi based on ER-stress, glycolysis and gliogenesis are highlighted on the UMAP.

#### Change of cellular identity in stressed cells

Besides increased stress-gene expression, stress clusters were characterized by low expression of pan-neural markers (*NEUROD6, DCX, MAP2, NCAM1, ELAVL4*; Appendix Figure S1E). To test if this marker depletion is also reflected by a general change in glial or neural fates, we extended our scoring approach. We calculated scores for the two cardinal cell states in neural development, neurogenesis (GO:0022008), and gliogenesis (GO:0042063). Both terms were depleted in stressed cells (**Fig 2A**). Compared to both glial and neural clusters, stressed cells also showed remarkably low scores for ‘cell differentiation’ (GO:0030154), and ‘forebrain development’ (GO:0030900), suggesting that stressed cells are in a more ‘basal’ state compared to differentiating neurons (Appendix Figure S2 AB).Thus, chronic stress in organoids comes at the expense of neurogenic cell differentiation and leads to a switch in cell fate. We therefore refer to the reduced expression of marker genes as the ‘stress identity’. As stressed cells lose both glial and neuronal identity and stop differentiating, it explains why these cells occupy the middle of the UMAP (**Fig 2A**).

#### Granular evaluation overcomes noise inherent to single-cell data

Single-cell gene expression measurements are inherently noisy. While GO-scores are computed across multiple genes per cell, these may still suffer from high variability and noise. Indeed, some cells within the stress-clusters showed low stress-scores, even if clustering together (**Fig 2A**). At the same time, sporadic cells in well-defined cell types showed high stress-scores. These cells expressed stress genes inconsistently, but expressed respective cell type markers, which are otherwise absent in stress-identity.

To overcome variability in single cell measurements, one can either denoise the data, e.g., by model-based imputations, or group cells and evaluate them together. Many different imputation methods have been developed recently, however, imputed values often vary (Hou et al., 2020), and they can induce false signals (Andrews and Hemberg, 2018). This is probably due to the complexity of the imputation problem. We therefore took a grouping approach where we partitioned cells into groups of 100-200 cells by ultra-high-resolution SNN-clustering in PCA-space (Methods) resulting in small groups of cells, that we term *granules*.

The ultra-high-resolution clustering approach can overcome the problems of boundaries by breaking down the data into minute groups of cells. To get sufficient coverage for robust gene scoring, and because clustering creates some very small granules, we added a reclassification step, where cells in granules with <30 cells are reassigned to the closest granule above threshold (Methods).

To test the granular approach, we compared stressed cells identified by Gruffi’s granular method (gSC) and stressed cells identified on single-cell scores (scSC). We compared cells only identified by either, both or neither of the approaches. First of all, scSC were evenly scattered across all clusters, while most gSC were close to stress-clusters and the cells identified by both methods (Appendix Figure S2C). By definition, single-cell selection on stress-scores identified the cells with highest stress-scores. However, scSC showed less of all other features defining stress-identity: lack of cell differentiation, lower mitochondrial and higher ribosomal mRNA content. Granular identification found cells that shared these features more with stress-identity (Appendix Figure S2D-I). We implemented both methods in Gruffi, but we concluded that the granular approach is more suitable if one aims to exclude stress-identity, whereas the single cell approach is more suitable if one aims to simply find cells with the highest stress gene expression, but otherwise properly specified cells.

#### Granular functional filtering (Gruffi) isolates and removes stressed cells

As clustering-based identification failed to detect stressed cells specifically and robustly, we built on the concepts above and developed Granular Functional Filtering or Gruffi (**Fig 2B**). Gruffi takes a number of Gene Ontology Pathways (1) to obtain corresponding gene sets (2), and computes cell-wise GO scores (2). At the same time, it identifies a suitable resolution (I), clusters cells in granules (II), and reassigns cells of too small granules (III). Merging these, it then computes the multiple granule scores (4), estimates a threshold separating stressed and non-stressed cells (5) and assigns a ‘stress’ label integrating multiple scores (6).

To uniformly determine the prevalence of stressed cells across organoids and protocols, we applied Gruffi to the integrated organoid dataset. After pathway scores calculation, we estimated the optimal clustering resolution, which was 1009 granules with a median of 154 cells (**Fig 2C**). Stress identification must be robust across all datasets, therefore we incorporated 3 scores: the two most specific pathways: ‘glycolytic process’ (GO:0006096) and ‘response to endoplasmic reticulum stress’ (GO:0034976), and a negative filter on ‘gliogenesis’ (GO:0042043) score accounting for the higher native expression of ER genes in glia. Addition of further negative filters did not improve identification; however, we implemented this option in our algorithm.

Gruffi then calculated the average and cell number normalized functional scores per granule (**Fig 2D**) resulting in a 3-dimensional functional score for each granule. This combinatorial approach is flexible to the type and number of scores used, which may be useful for applications beyond its original scope. Next, Gruffi estimated the thresholds separating stressed from normal cells, accounting for score variability among non-stressed cells (Methods). At this point the retrieved thresholds can and are advised to be inspected and refined via the implemented interactive Shiny App interface. Throughout our analysis presented here, we did not further adjust the by Gruffi initially estimated thresholds. Finally, combining all thresholds, Gruffi classified stressed and non-stressed cells (**Fig 2F**), which largely overlapped with the expression of key stress markers, and with high stress scores (**Fig 1C**). While this suggested a correct identification, we continued the in-depth analysis of stressed cells.

### Stress has a profound, yet limited transcriptional effect and stress-identity is not present in vivo

#### Stressed cells show a profound transcriptomic change

As all samples had stress-identity cells, we searched for their defining features and their consequences on the whole organoid. Stressed cells fell in two distinct clusters with either a more glial or a more neural character (**Fig 1**). We therefore divided Gruffi’s stressed cells into these two categories to investigate the heterogeneity of stress response in organoids (methods). We quantified the total expression of mitochondrially encoded genes and of ribosomal mRNAs, which correspond respectively to ∼2% and ∼10% of the total transcriptome, respectively. These are widely used to assess quality and cell state in scRNAseq (Luecken and Theis, 2019). We hypothesized that chronic hypoxia and glycolysis diminish the need for oxidative phosphorylation, which may translate to fewer mitochondrial UMIs. Indeed, stressed neurons showed 52%, whereas stressed progenitors showed a 25% reduction in mitochondrial read fractions, as compared to their non-stressed counterparts (**Fig 3A**). In contrast, ribosomal mRNA fractions were 40% and 23% higher, respectively (**Table S2**), perhaps to compensate for ER-dysfunction (MWW p.val < 2e-16 in all cases)

**Fig. 3:**
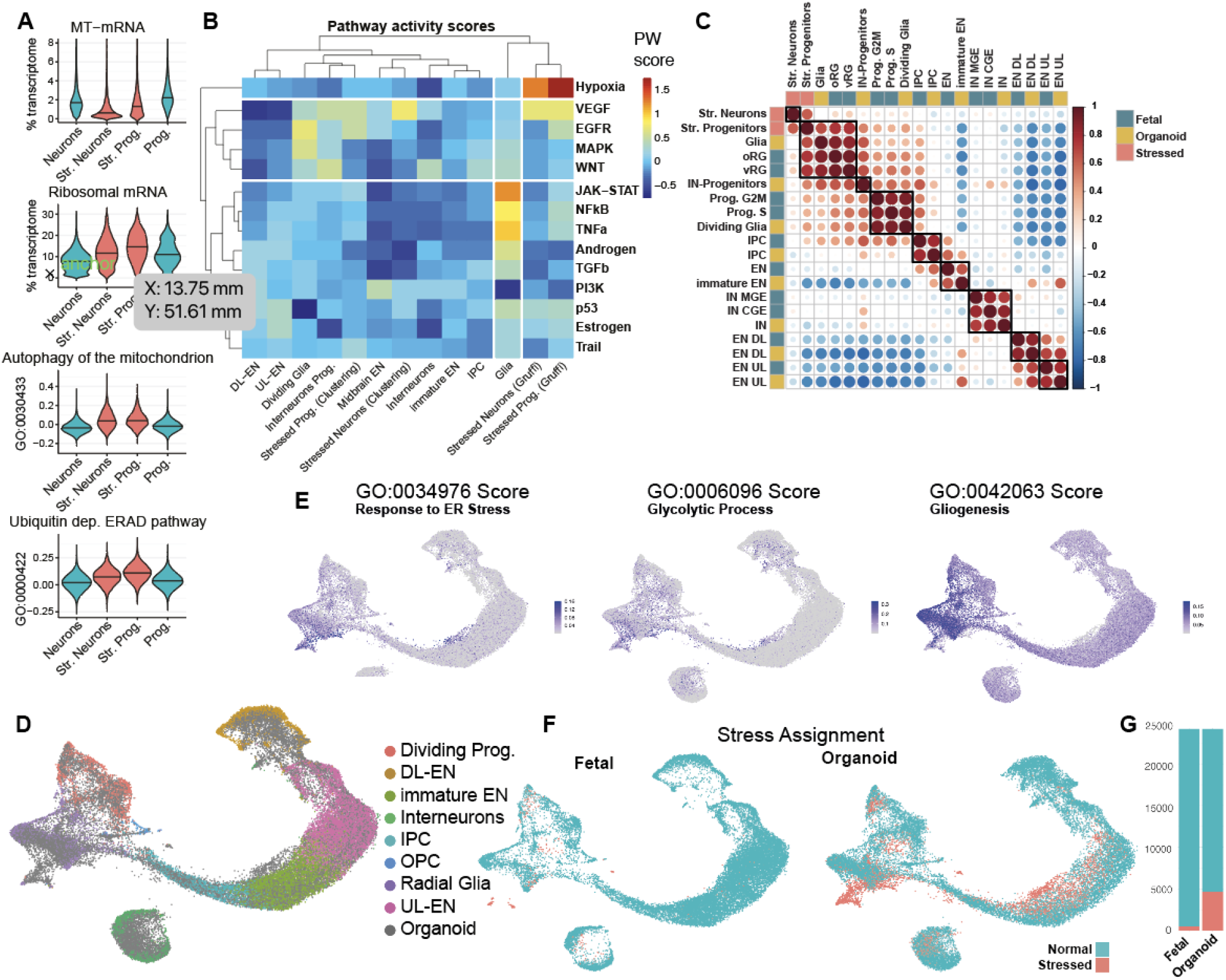
Stress has a profound, yet limited transcriptional effect, and stress-identity is not present in vivo. (**A**) Functional consequences of stress on the transcriptome. Mitochondrial (MT) mRNAs are decreased (top), while ribosomal mRNA are increased in stressed neurons and progenitors (second from top). Mitochondrial autophagy is increased in stressed cells, possibly explaining the reduced MT-mRNA (third from top). increased protein degradation via the ERAD pathway in the ER is a likely reason for increased ribosomal reads. (**B**) Activity of multiple pathways defined by PROGENy clearly separates stressed cells. Hypoxia is the only uniquely activated pathway in stressed cells. Hierarchical clustering based on pathways independently validates that stressed clusters as identified by Gruffi are distinct from other cell types. (**C**) Correlation of fetal reference to organoid clusters. Only clusters that are found in both datasets and stressed cells are shown. Gene modules of co-regulated genes were computed in the fetal reference data. Aggregated expression per cluster and module was correlated. The color code marks the origin of the clusters (blue for fetal and yellow for organoid). Stressed neuron and progenitor clusters are marked in orange. Fetal clusters correspond to original clusters with adjusted names (Polioudakis et al., 2019). (**D**) CCA Integration of ∼24K fetal cells and an equal number of randomly sampled organoid cells from Fig 1B show that most cell types intermingle. Cluster annotation represents the original annotation of the fetal dataset. Gray points represent organoid cells. (**E**) Gruffi’s single-cell pathway scores for ER-stress (GO:0034976), glycolytic process (GO:0006096) and gliogenesis (GO:0042063) on UMAP. Granule clustering at resolution 37 (determined by Gruffi), resulting in 249 granules with a median of 193 cells per granule, after reassignment of small clusters. (**F**) Stress cell assignment by Gruffi identifies the vast majority of stressed cells in organoid samples. (**G**) Quantification of F. In total 5171 cells (10,68 % of all cells) were identified as stressed. 523 of these are fetal (2,16% of fetal cells) and 4648 cells are from the organoid datasets (19,2% of organoid cells).

#### Increased catabolism in stressed cells

Because we saw the decrease in mitochondrial mRNAs, we looked for transcriptomic signatures for the active degradation of mitochondria. To that end, we applied Gruffi’s scoring method for relevant GO terms, and found opposite signs: both stressed groups showed positive scores for ‘autophagy of the mitochondrion’ (GO:0000422) on average, while normal cells do not (**Fig 3A**). At the same time the groups scored similarly for ‘autophagy’ (GO:0006914) (**Appendix Figure S3A, Table S2**).

Translation in ER stressed cells might lead to protein degradation via the ubiquitin-dependent ERAD pathway. Therefore, we calculated the corresponding score (GO:0030433), and as for mitophagy, found that stressed cells scored positively, while normal cells scored negatively (**Fig 3A, Table S2**). Altogether, these changes show that stress induces major changes to the cell’s physiology and metabolism that go beyond acute stress response.

Finally, we analyzed cell types by clustering PROGENy’s activity-score across signaling pathways and clusters (Schubert et al., 2018). We found that stressed cells form an outgroup and are marked by the upregulation of Hypoxia and VEGF pathways, and the downregulation of the PI3K pathway, highlighting oxygen deficiency and quiescence (**Fig 3B**, methods). To ask how stressed cells identified by Gruffi differ from those identified by the naïve approach of identifying stress by clustering (**Fig 1**), we subcategorized stressed cells identified in only one of the two classifications. The complete lack of separation of cells found by the naïve approach contrasted the salient stress features found by Gruffi (**Fig 3B**).

#### The presence of stressed cells does not affect specification and maturation of non-stressed neurons^1^

A previous study reported that stress in organoids leads to impaired cell-type fidelity, and incomplete maturation as a global phenomenon in organoids (Bhaduri et al., 2020). To test those effects, we compared cell types in the organoid datasets to fetal cortical data of comparable age (de la Torre-Ubieta et al., 2018; Polioudakis et al., 2019) (methods). We defined the fetal brain as the reference data and constructed modules from co-expressed genes (**Table S3**, methods) as described before (Trapnell et al., 2014). The resulting 65 aggregated gene modules were then used to calculate Pearson correlation across clusters and then visualized in a heatmap (**Fig 3C**). Major cell types (excitatory neurons, interneurons, and progenitors) formed the largest clusters across the dataset. Most organoid cell types pairwise best matched the corresponding fetal cell type, indicating that organoids undergo proper cell type specification unlike suggested previously (Bhaduri et al., 2020). Stressed neurons, however, were uncorrelated to all cell types, except stressed progenitors (**Fig 3C**). This dissimilarity to all fetal and organoid cell types, along with the analyses above, indicated that stress neurons lost most of their identity and formed a new transcriptional cell-state. Interestingly, while stressed progenitors showed a similarity to stressed neurons, they also showed a strong progenitor identity, suggesting either an increased robustness or more distinct transcriptome of the progenitor state. Altogether we found no evidence of imparied cell-type fidelity in organoids, as cell types in organoids match respective cell types *in vivo*, and that stressed cells show little resemblance to cell types found in the fetal cortex.

#### Stressed cells in organoids have no fetal counterpart

Stressed cells might also exist in vivo, even if they did not form a recognized cluster in published studies. While some previous reports have argued that stressed cells similar to organoids exist *in vivo* (Gordon et al., 2021; Tanaka et al., 2020), others claim that it is an artifactual population specific to organoids (Bhaduri et al., 2020). Therefore, we integrated the fetal brain dataset with a matching number of randomly downsampled cells from the organoid dataset. Using the published fetal cell type annotation, we found that the organoid dataset was generally well recapitulating the fetal data (**Fig 3D**). At the same time, CGE and MGE (caudal and medial ganglionic eminences) interneuron differences and deep layer excitatory neuron differentiation were clearer in fetal data.

Interestingly, there were two populations entirely of organoid origin: a population near the neural trajectory (I) and a progenitor population (II, **Fig 3D**). To test for stress identity, we calculated stress scores as before, and found that precisely these populations score high for ER stress and glycolysis (**Fig 3E**). To quantify stressed cells and validate our method on both datasets, we ran Gruffi on the integrated object. This identified stressed cells almost exclusively in the two aforementioned populations (**Fig 3G**) and almost exclusively in organoid samples (**Fig 3H**). Altogether, our analysis suggests that stress-identity is an in vitro artifact and there is minimal to no stress-identity *in vivo*.

### Stressed cells do not affect the maturation of other cells, but their removal improves data quality and interpretability

#### The removal of stressed cells reveals clear trajectories that recapitulate fetal neurodevelopment

Removing stressed cells might create a better model of the fetal brain development. Stress genes majorly contribute to variable genes, which determine both clustering and visualization (Luecken and Theis, 2019). We therefore removed all stressed and low-quality cells, reidentified variable genes, and recalculated all dependent representations (PCA, UMAP, snn-graph) and downstream analysis with identical parameters.

Starting from the progenitors, the resulting UMAP revealed 3 trajectories (“E”, “I” & “MB” in **Fig 4. A-B**), representing cortical excitatory, cortical inhibitory, and midbrain neurons, respectively. Before removing stressed cells, no midbrain trajectory was visible although midbrain cells were obviously present (**Fig 1B**). Instead, midbrain cells were linked to their progenitors only by a small, separated population of the ‘yellow’ cluster. After stressed cell removal, these trajectories now lead to distinct populations of mature neurons, as opposed to the continuum of connected clusters before Gruffi (before stress removal). Notably, this lineage separation recapitulates fetal neurodevelopment.

**Fig. 4:**
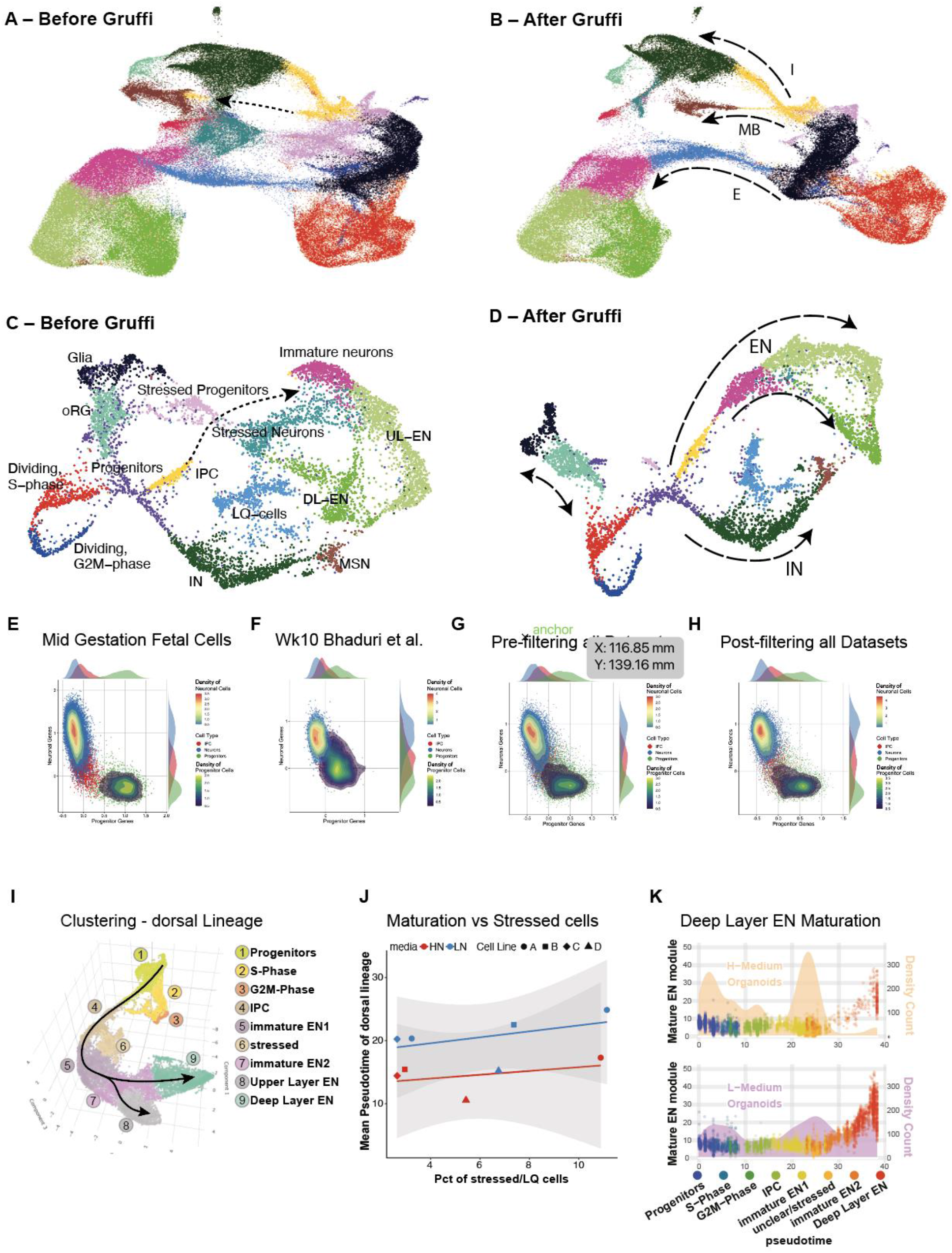
Stressed cells do not affect the maturation of other cells, but their removal improves data quality and interpretability. Lineage trajectories improve with Gruffi. (**A-B**) UMAPs of the integrated organoid dataset before Gruffi (**A**), and after Gruffi (**B**). Dotted arrow in (A) highlights the broken trajectory in the development of midbrain cells. Dashed arrows in (B) highlight the developmental trajectories of interneurons, midbrain neurons, and cortical excitatory neurons. Clustering and colors are the same as in (Fig 1A). (**C-D**) We repeated the analysis on a smaller subset of the data downsampled to ∼8200 cells, a typical outcome of a single 10X experiment (this subset does not contain midbrain neurons). Note that almost any lineage relationship could be inferred from the representation before Gruffi (A), but not that excitatory neurons relate to IPCs (dotted line). After Gruffi (B) the lineage trajectories show the known relationships allowing, for instance, pseudotime calculations. (**E**) Progenitor- (x-axis) vs. excitatory neuron- (y-axis) scores on mid-gestational fetal cells (ref Geschwind, **Fig EV4B**) identifies two separate populations. Each cell is colored based on neuronal or progenitor cell cluster identity. Color depicts progenitors (green), intermediate progenitor cells (IPC, red) and neurons (blue). Additionally, the density of progenitors (blue to yellow) and neurons (blue to red) is shown. The margins of the plot depict the density distribution of the three different cell types across the progenitor (x-axis, top) and neuron (y-axis, right) score. (**F**) Example dataset showing impaired cell subtype specification, as described in a previous study (ref Bhaduri). (**G**) Subtype specification of all datasets analyzed in this study. While most neurons and progenitors show high values of the respective scores, there are some cells without specification to either neurons or progenitors (see **Fig EV4C-F** for individual datasets). (**H**) Subtype specification after filtering out stressed cells. The remaining cells specify properly to neurons and progenitors, with only IPCs localizing between the two populations (see **Fig E4C-F** for individual datasets). (**I**) Clustering of dorsal lineage cells of datasets grown in two separate media conditions (materials and methods). Pseudotime analysis was performed from progenitors (cluster 1) to upper layer (cluster 8) and deep layer neurons (cluster 9, see **Fig EV4G** for pseudotime plot). (**J**) The maturation along pseudotime is measured as the mean pseudotime value of neurons (y-axis) and plotted against the percentage of stressed cells identified in the datasets (x-axis). The color code shows the two different media formulations of individual datasets (marked by different symbols) and the linear regression model (line with CI). The percentage of stressed cells did not correlate with the maturation difference of non-stressed neurons. (**K**) Maturation of deep layer excitatory neurons (DL-EN) in two different media formulations. Cells are color coded for clusters (**Fig 4I**) and plotted along pseudotime (x-axis). The density-count of cells along pseudotime is shown behind the dots (yellow area for H-medium, purple area for L-medium, right y-axis). The position of the points reflects the expression of a module of co-regulated genes enriched in DL-EN (left y-axis, **Fig EV4H**). While proper maturation indicated by expression of the DL-EN module occurs in both media conditions, in L-medium maturation is much more frequent (see **Fig EV4I-K** for upper layer ENs and for individual datasets).

To ensure the robustness of our approach we repeated the analysis on a single dataset (**Fig EV2A** to **E**), as well as downsampling the full dataset to ∼8200 cells, which is the typical output of a single 10X experiment (**Fig 4C-D**). Both analyses revealed that expected lineage trajectories are missing or broken in the UMAPs before Gruffi (midbrain in **Fig 4A**, and excitatory neurons in **Fig 4C**), but they are correctly recovered after running Gruffi (B, D). Correct and continuous trajectories in low dimensional representations are essential for most pseudotime methods that use these as basis for pseudotemporal cell assignment.

We then tested if our method can be applied to independent datasets by re-analyzing the data from a recent publication reporting a large “undefined” cluster (Samarasinghe et al., 2021). We ran Gruffi on the precomputed R data object obtained from the authors and marked stressed cells (**Fig EV3A** and **B**). This dataset is particularly well suited to demonstrate the versatility of Gruffi, as it contained secretory choroid plexus cells, which are undergoing high physiological ER-stress due to secretion. We therefore derived a choroid plexus score as an additional negative filter score. As no GO-term exists for choroid plexus, we turned to the recent publication of choroid plexus organoids (Pellegrini et al., 2020) identified marker genes, and derived the choroid plexus score (methods). Gruffi labeled 74% of the “Undefined” cluster as stressed and an average of 1% of cells from other clusters (**Table S4**). We then performed DGEA on stressed vs. non-stressed cell and GO-term enrichment on all enriched coding genes (methods). Visualization of enrichments on the protein interaction network using STRING (**Fig EV3C**) showed that apoptosis, stress, unfolded protein response, and hypoxia dominate cells identified as stressed in this dataset as well.

#### Organoids show proper cell type specification along the excitatory lineage

A previous study reported that stress in organoids not only leads to impaired cell-type fidelity but also incomplete maturation (Bhaduri et al., 2020). To investigate whether stressed-identity cells affect specification along the excitatory lineage, we compared progenitor and neuron signatures (methods, **Table S5**). This led to the expected bimodal separation of the fetal samples (**Fig EV4A**). As this dataset was used to derive the signatures, we confirmed the separation of neurons vs progenitors using an independent fetal dataset with multiple samples around mid-gestation (**Fig 4C, Fig EV4B**). Next, we asked how organoid samples separate using those signatures. We calculated signature scores on Bhaduri et al. organoid datasets and reproduced the previously reported lack of specification (**Fig 4D**). In contrast, when testing the other datasets analyzed in this study, we detected a fetal-like specification (**Fig 4E, Fig EV4C-F**), suggesting proper specification in most organoid datasets.

Nevertheless, organoid cells still separated less than fetal cells, and more cells were scoring low on both progenitor and neural axes. As stressed cells are characterized by the lack of both glial and neural signatures (**Fig 2A**), we hypothesized that stressed cells may populate the area between neurons and progenitors. After annotating stressed cells, we found two populations in between progenitors and neurons: stressed cells and intermediate progenitors, or IPCs. After stress removal, most of those remaining cells are IPCs (**Fig 4F**, in red), which are indeed a transitory stage between glia and neurons. Altogether, we find no evidence for general misspecification in organoids. Instead, progenitors and excitatory neurons properly separate, while two specific populations, IPCs, and stressed cells, lack specific neuronal or progenitor signatures.

#### The presence of stressed cells does not affect the maturation of other cell types

Besides a lack of cell type specification, incomplete maturation due to stress was previously suggested (Bhaduri et al., 2020). As a positive control for maturation, we took two media conditions that affected maturation (Eichmüller et al., 2022). We grew pairwise matched samples in two different media conditions, then analyzed together (**Fig 4G**). We calculated the pseudo-temporal trajectory of the excitatory lineage and graded each dataset for maturation along this trajectory (**Fig EV4G**, methods).

Our results showed that the fraction of stressed cells does not explain maturation differences (**Fig 4H**), but the choice of media does: low-nutrient media improved neural maturation. To understand the impact on maturation on single cell level, we plotted individual cells along the maturation trajectory, split by media condition (**Fig 4I**). Additionally, to assess expression changes associated with mature states we generate scores for mature neurons (**Fig 4I, Fig EV4H** to **J, Table S6**). This revealed that while organoids grown in either condition had abundant excitatory neurons, those in HN media remained mostly immature, while organoids grown in LN media contained more mature neurons (**Fig 4I, Fig EV4J** to **L**). In sum, the presence and abundance of stressed cells in a sample have negligible effects on neural maturation, while measurable differences arise by the choice of media.

## Discussion

Brain organoids generate complex, structured tissue *in vitro* (Eiraku et al., 2008; Kadoshima et al., 2013; Lancaster et al., 2013; Paşca et al., 2015; Qian et al., 2016). Besides their tremendous potential for modeling human diseases (Sidhaye and Knoblich, 2021), it is critical to understand and account for their limitations. Here, we showed that a population of stressed cells exists in all analyzed organoid samples, and that this is a biologically distinct population, which is not found *in vivo*. We provide an in-depth analysis of these cells that hopefully will help to decipher the needs of 3D tissue in culture.

While an experimental solution is the end goal, stress is present in published and current experiments. To tackle this issue, we developed Gruffi, an *in silico* tool to bioinformatically identify and remove these cells, based on stress pathway activity scoring. Gruffi comes with a graphical and interactive interface. It integrates into a typical single-cell analysis workflow using Seurat, but can be used in other pipelines as well. The resulting stress-decontaminated samples displayed a clearer representation of the fetal neural development and showed higher similarity to *in vivo* samples. Even if future organoid protocols may resolve cellular stress, earlier published data still suffers from stress, which negatively impacts data integration. Gruffi, however, can recover such data for comparison, reducing the need for performing new experiments.

We observed diverse stress pathway activity, and it is important to understand how they are connected on a cell biological level. Our results are compatible with earlier observations that the organoid core, but not surface, is hypoxic (Qian et al., 2019), explaining why stress characterizes only in a defined set of cells. The central role of hypoxia can explain the other transcriptional shifts. The lack of oxygen triggers a metabolic shift, from oxidative phosphorylation to anaerobic glycolysis.

Hypoxia also triggers ER stress, in two ways. First, glycolysis is much less efficient in energy production, leading to energy depletion, and consequently stronger metabolite transport is needed. These transporters are secreted via the ER-pathway (Loike et al., 1992), triggering unfolded protein response (UPR) (Lee et al., 2020). At the same time, the depletion of energy leads to a pH imbalance, affecting organelles that rely on ATP-dependent transporters for ion homeostasis (Chiche et al., 2010).

Our results are consistent with a previous observation that acute hypoxia in cortical spheroids triggers a strong ER-stress response (Pa?ca et al., 2019). However, a simplistic, one-dimensional distance-to-surface model of nutrient availability cannot explain the heterogeneity of stress marker expression. It is an interesting future direction to determine different cellular niches, e.g. by local variation in oxygen and nutrient levels. Similarly, an interesting question for future studies is, how cellular heterogeneity leads to the differential expression of stress markers in close neighboring cells.

Importantly, stress identification is just the first application and proof of principle for granular functional filtering. This flexible framework can be extended to many other applications in single-cell analysis. As long as a group of cells form an identity, so that they group together in any low dimensional representation, and coexpress a defined geneset (GO-terms, KEGG-pathways, etc.), the cells can be identified, studied and removed. Here, we applied Gruffi to remove stressed cells from brain organoid datasets, but we think that there are many other applications possible, such as selecting cells from a lineage or cells responding to a treatment. Currently, brain organoids are the largest and longest-cultured 3D organoid systems and are therefore particularly affected by stress. As 3D tissue models and investigations become ever more sophisticated, cell culture induced artifacts are more important to account for. Therefore, we expect that our approach will find many applications beyond its original scope.

## Supporting information

All supplementary tables

## Data and code availability

The single-cell RNA-sequencing data have been uploaded to Gene Expression Omnibus (GEO) under reference number GSE1XXXXXXXXX (ncbi.nlm.nih.gov/geo/query/acc.cgi?acc= GSE1XXXXXXXXX). We used publicly available raw sequencing data from the following publications (Bhaduri et al., 2020; Eichmüller et al., 2022; Kanton et al., 2019; Polioudakis et al., 2019, 2019; Velasco et al., 2019) and obtained wild type patient data from the authors of, complying with ethical and data safety requirements (Khan et al., 2020; Samarasinghe et al., 2021). The Gruffi package will be available under github.com/jn-goe/gruffi. The code for analysis will be accessible on Github: github.com/vertesy/Limited.Stress.in.Brain.organoids. The following custom function libraries were used for the analysis: *Stringendo, ReadWriter, CodeAndRoll2, MarkdownHelpers, ggExpress, Seurat*.*Utils*, all freely available under github.com/vertesy.

## Acknowledgements

We thank all Knoblich laboratory members for continued support and discussions. We thank Ilaria Morassut for helping with preparing some of the sequencing libraries and Angela M Peer for growing some of the organoids. We thank Lina Dobnikar and Thomas Krausgruber at CeMM for assistance with generating and interpreting some of the sequencing data. We thank Jessie Buth, Ranmal Samarsinghe and Ben Novitch for providing processed data and help with (Samarsinghe et al., 2021). We thank the IMP/IMBA and Vienna Biocenter Core Facilities (VBCF) for their valuable expertise and committed work, in particular Gerald Schmauss at the IMP/IMBA biooptics facility and the IMP/IMBA Molecular Biology Service. Abel Vertesy received funding from EMBO LTF:11122019. Work in A.v.H.’s laboratory is supported by the Austrian Science Fund (project SFB-F78 P11). Work in J.A.K.’s laboratory is supported by the Austrian Federal Ministry of Education, Science and Research, the Austrian Academy of Sciences, the City of Vienna, a Research Program of the Austrian Science Fund FWF (SFBF78 Stem Cell, F 7803-B) and a European Research Council (ERC) Advanced Grant under the European 20 Union’s Horizon 2020 program (no. 695642) and the European Union’s Horizon 2020 research and innovation programme funded project HCA|Organoid (no. 874769). JN is supported by the Austrian Science Fund (FWF) project number F78 to Arndt von Haeseler.

## Author Contribution

AV, OLE, and JAK conceived and JAK and AH supervised the project. AV and JN developed the stress identification algorithm. Data analysis was performed by AV, OLE, JN, MN. Organoids were grown by CE, MBE and sequencing libraries were prepared by AV, SL. AV and OLE wrote the manuscript with input and editing from JAK and JN. Funding Acquisition: JAK, AvH, CB, AV.

## Figure Legends

Main Figure Legends (see separate files for higher resolution)

## Extended View Figure Legends

**Fig EV1.**
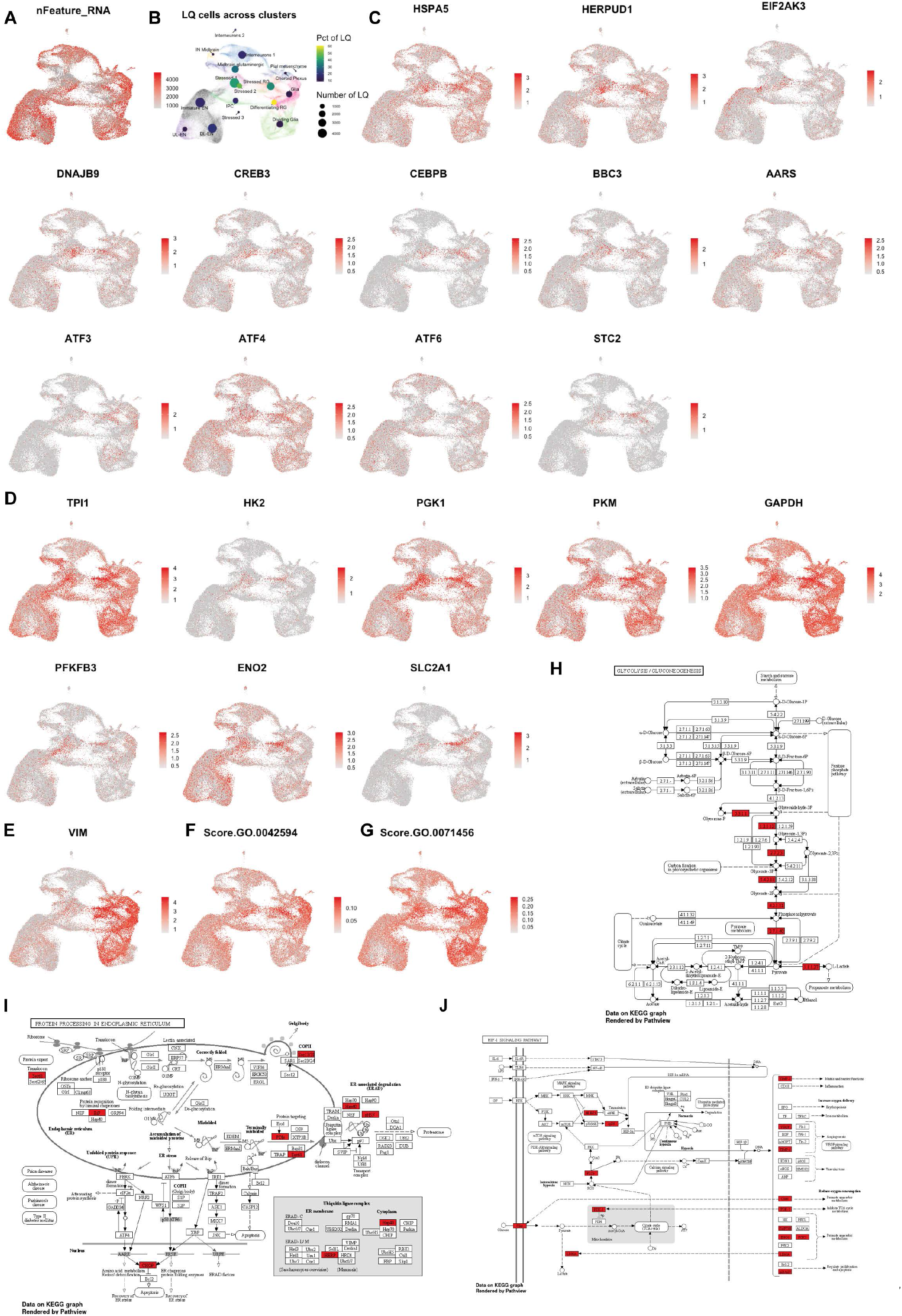
Metabolic changes and marker genes is stressed cells. (**A**) UMAP of organoid integration shown in Fig 1B colored by number of RNA features per cell (nFeature_RNA). (**B**) Low-quality (LQ) cells as determined by expression of less than 1000 features. In the background the clustering of Fig 1B is shown. On top per cluster the percentage of LQ cells per cluster (color) and the number of LQ cells (size) are depicted. (**C**) Expression of additional endoplasmic reticulum (ER) stress genes enriched in the stress clusters. (**D**) Expression of additional glycolysis genes enriched in the stress clusters. (**E**) Vimentin (*VIM*) is expressed in all progenitor populations regardless of lineage or stress state. (**F**-**G**) Additional GO-terms scores ‘response to starvation’ (GO:0042594) and ‘cellular response to hypoxia’ (GO:0071456) are also characteristic of stressed cells. (**H-J**) Stress cluster marker genes in relevant significantly associated KEGG pathways: HIF-1α signaling, (Genes: 12, Fold Enrichment: 14.4, FDR: 2.30e-9); Glycolysis, (7, 13.7, 2.3e-5); Protein processing in the ER, (8, 5.7, 1.9e-3). The top 150 coding stress marker genes were used for this analysis (as in Fig 1E). Enriched genes are marked red.

**Fig EV2.**
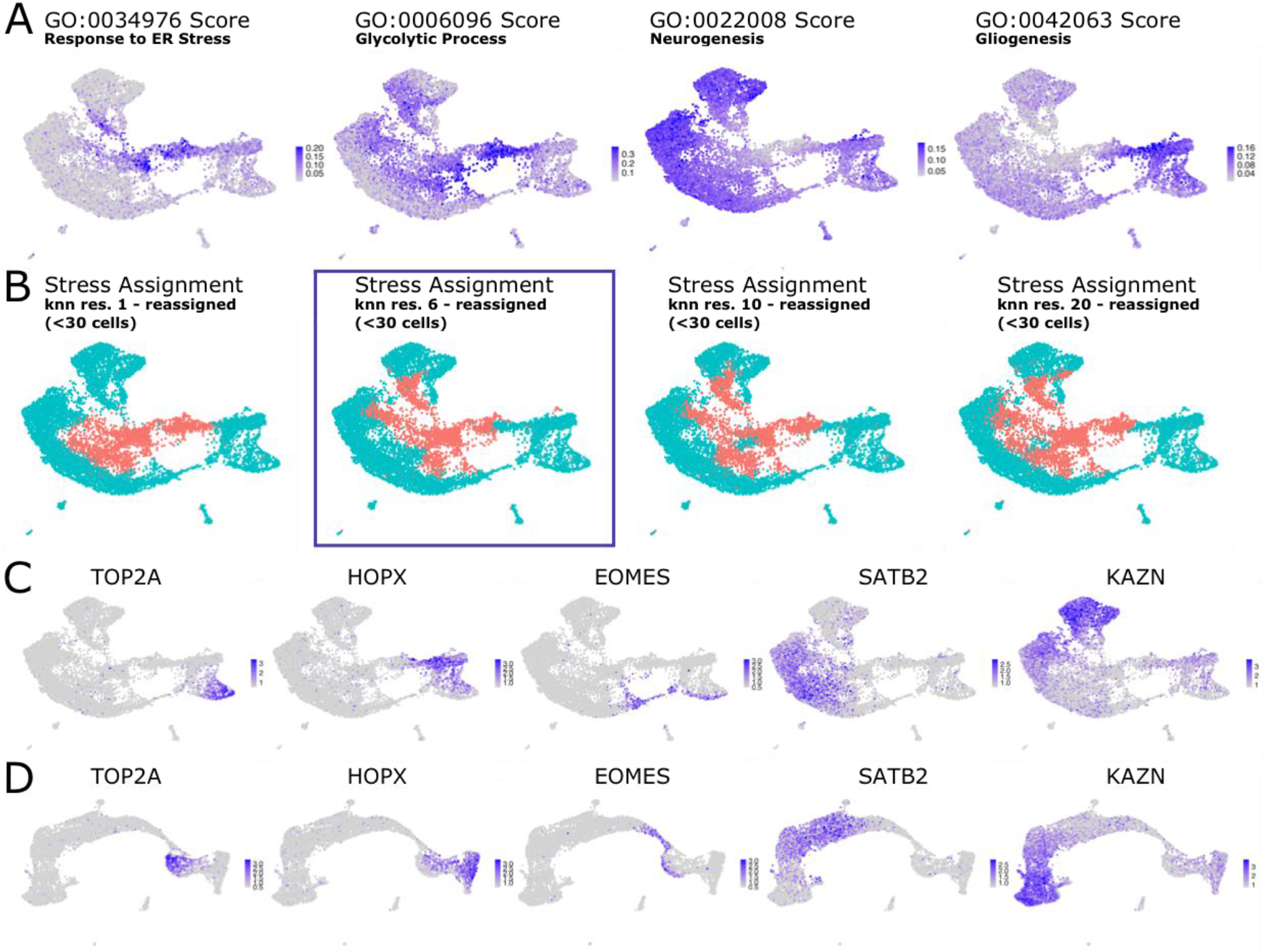
Benchmarking Gruffi on a single-experiment sized dataset. (**A**) UMAP plots of one dataset (Velasco 7) cell wise GO score for response to Endoplasmic Reticulum stress (GO:0034976), glycolytic process (GO:0006096) and gliogenesis (GO:0042063). (**B**) Stress assignment performed on k-nearest-neighbor clustering with resolution 1, 6, 10 and 20 plus reassignment of granules with a cell count below 30 (left to right). Resolution 6 (+ reassignment, the proposed resolution for a median cell number between 100 and 200 cells, see Methods), resulted in 53 granules with a median cell number of 197. (**C**) and (**D**) Expression profiles of markers for progenitor cells (*TOP2A, HOPX*), Intermediate Progenitors (*EOMES*), upper layer excitatory neurons (*SATB*) and deep layer excitatory neurons (*KAZN*) show that the developmental trajectory is refined in a newly computed UMAP after stress filtering (**D**) compared to before stress filtering (**C**). For the new UMAP, we recomputed (and scaled) the most variable genes, Principal Components and the UMAP embedding.

**Fig EV3.**
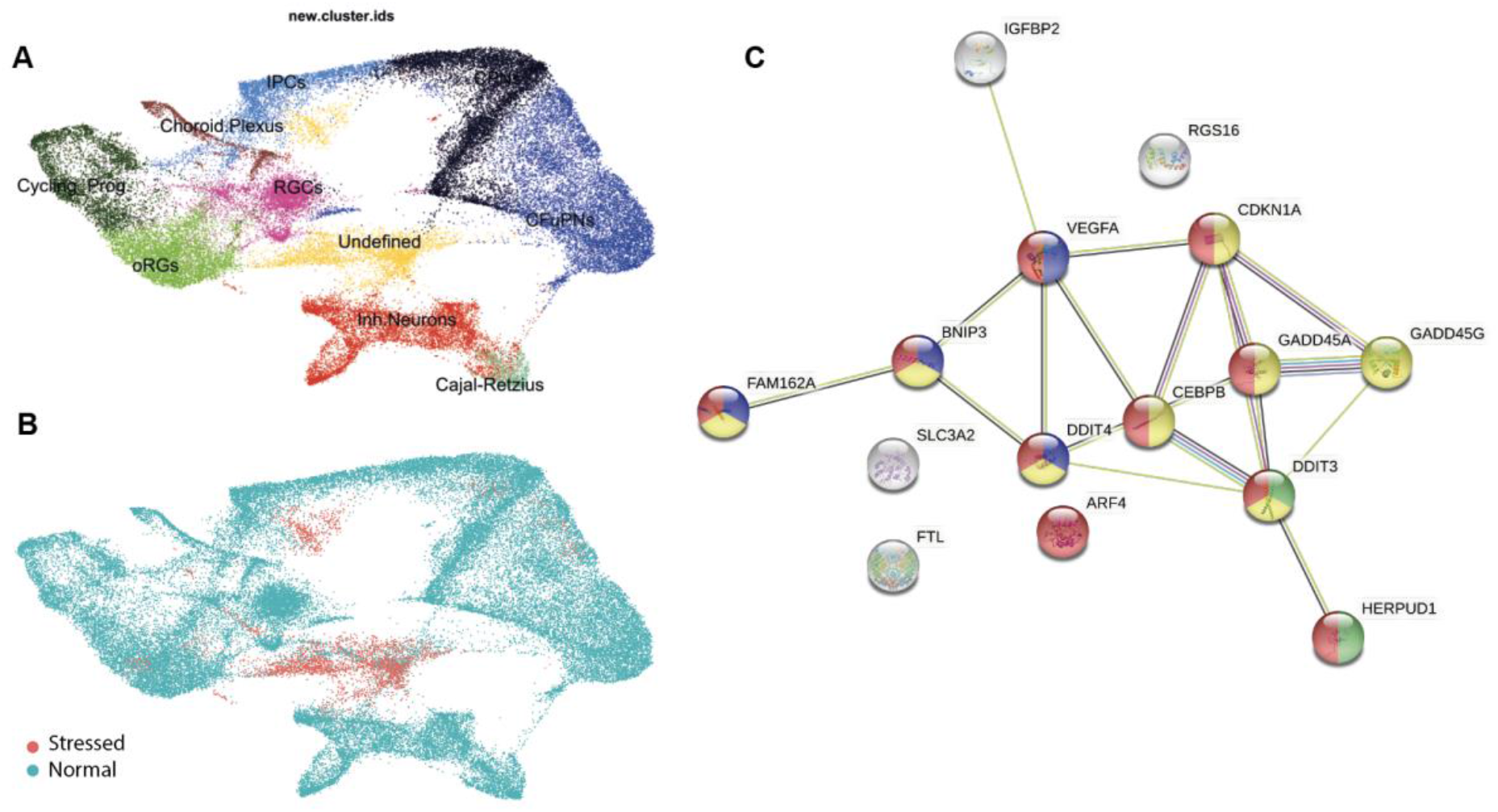
Stress identification in Samarasinghe et al. 2021. (**A**) Reproduction of Fig 4A UMAP in Samarasinghe 2021. (**B**) Stress assignment by Gruffi using res.25 (auto determined resolution, 260 granules), red is stressed, turquoise is unaffected. (**C**) Protein interaction map of marker genes of stress cells. Response to stress (red, GO:0006950, 0.02); Apoptotic process (yellow, GO:0006915, FDR=0.0003); PERK-mediated unfolded protein response (limegreen, GO:0036499, 0.0221); Response to hypoxia (blue, GO:0001666, 0.0334); DGEA and all enrichment terms are in (Table S6).

**Fig EV4.**
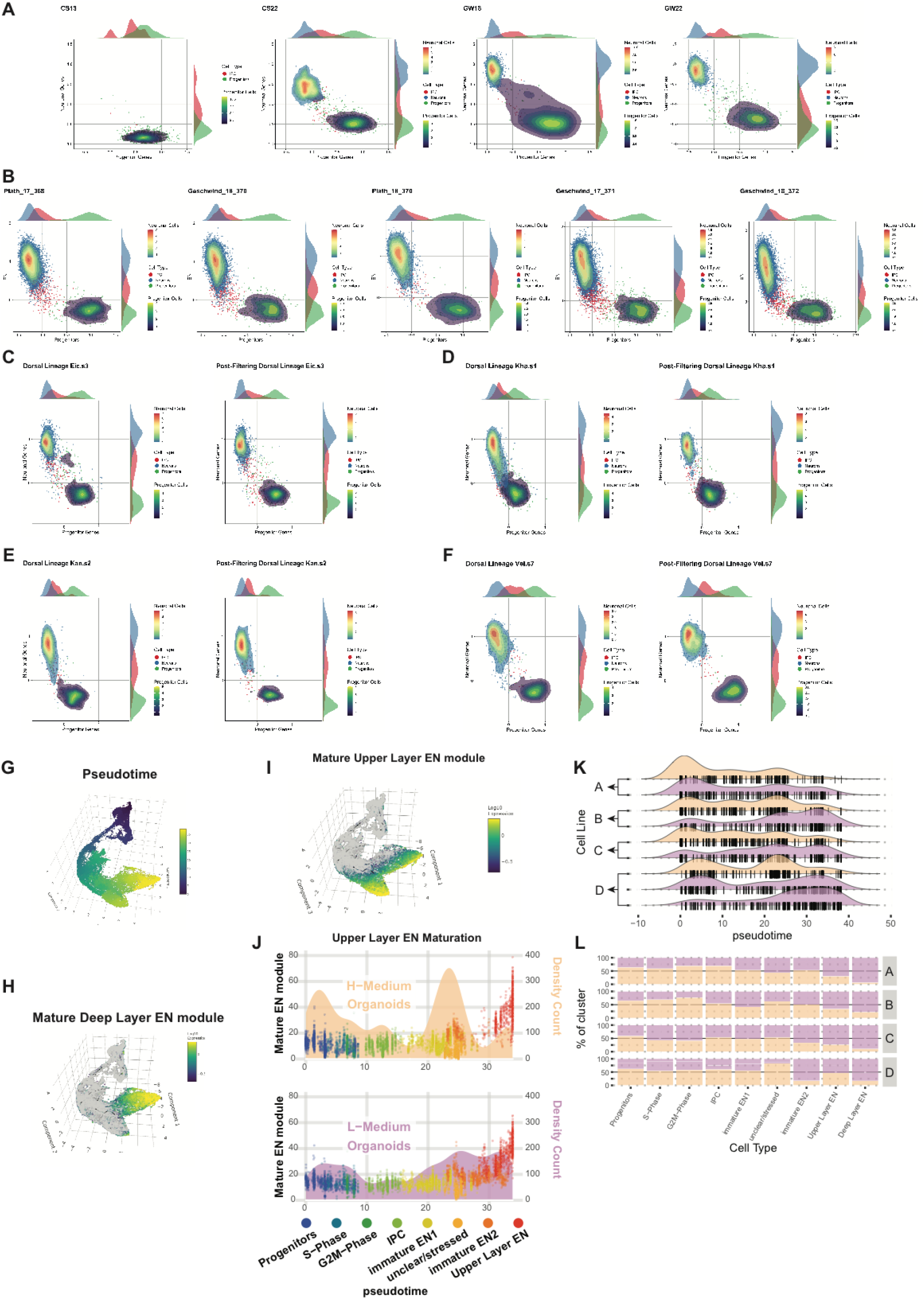
Proper specification and maturation in non-stressed cells. (**A**) Progenitor (x-axis) and excitatory neuron scores (y-axis) calculated in fetal brain datasets across brain development (materials and methods, Bhaduri et al 2020). Early datasets from Carnegie stage 13 (CS13) are enriched in progenitors, while late datasets from gestational week (GW) 18 and 22 show both neurons and progenitors. (**B**) Progenitor (x-axis) and excitatory neuron scores (y-axis) applied to multiple datasets of mid-gestation (**Fig 4E**) show reliable separation of neurons and progenitors, while only intermediate progenitors (IPCs, red) cluster in between the two populations. (**C**) to (**F**) Examples of pre- and post-filtering plots for subtype specification of individual datasets analyzed in this study. After filtering out stressed cells only IPCs remain in between neurons and progenitors. (**G**) 3D UMAP of dorsal lineage also shown in **Fig 4I** colored for pseudotime. (**H**) Expression of the deep layer excitatory neuron (DL-EN) gene module (Table S7) is specifically enriched in the DL-EN cluster (Cluster 9 in Fig 4I). (**I**) Expression of the upper layer excitatory neuron (UL-EN) gene module (Table S7) is specifically enriched in the UL-EN cluster (Cluster 8 in Fig 4I). (**J**) Maturation of UL-EN in two different media formulations analogous to DL-EN maturation in **Fig 4K**. Cells are color coded for clusters (**Fig 4I**) and plotted along pseudotime (x-axis). The density-count of cells along pseudotime is shown behind the dots (yellow area for H-medium, purple area for L-medium, right y-axis). The position of the points reflects the expression of a module of co-regulated genes enriched in UL-EN (left y-axis, **Fig EV4I**). While proper maturation indicated by expression of the UL-EN module occurs in both media conditions, in L-medium maturation is much more frequent. (**K**) Distribution of cells (black lines) and densities (areas) of individual organoids across pseudotime. Organoids derived from the same cell line (A to D) grown in the different media conditions (yellow for H-medium, purple for L-medium) are shown on top of each other. (**L**) Cluster contributions per individual organoids (as shown in **Fig EV4K**). The increased maturation in L-medium organoids is also reflected by higher proportions of mature cell types (UL- and DL-EN) in L-medium organoids. Cell numbers were downsampled to account for different library sizes.

## Supplementary Figure Legends

**Appendix Figure S1.**
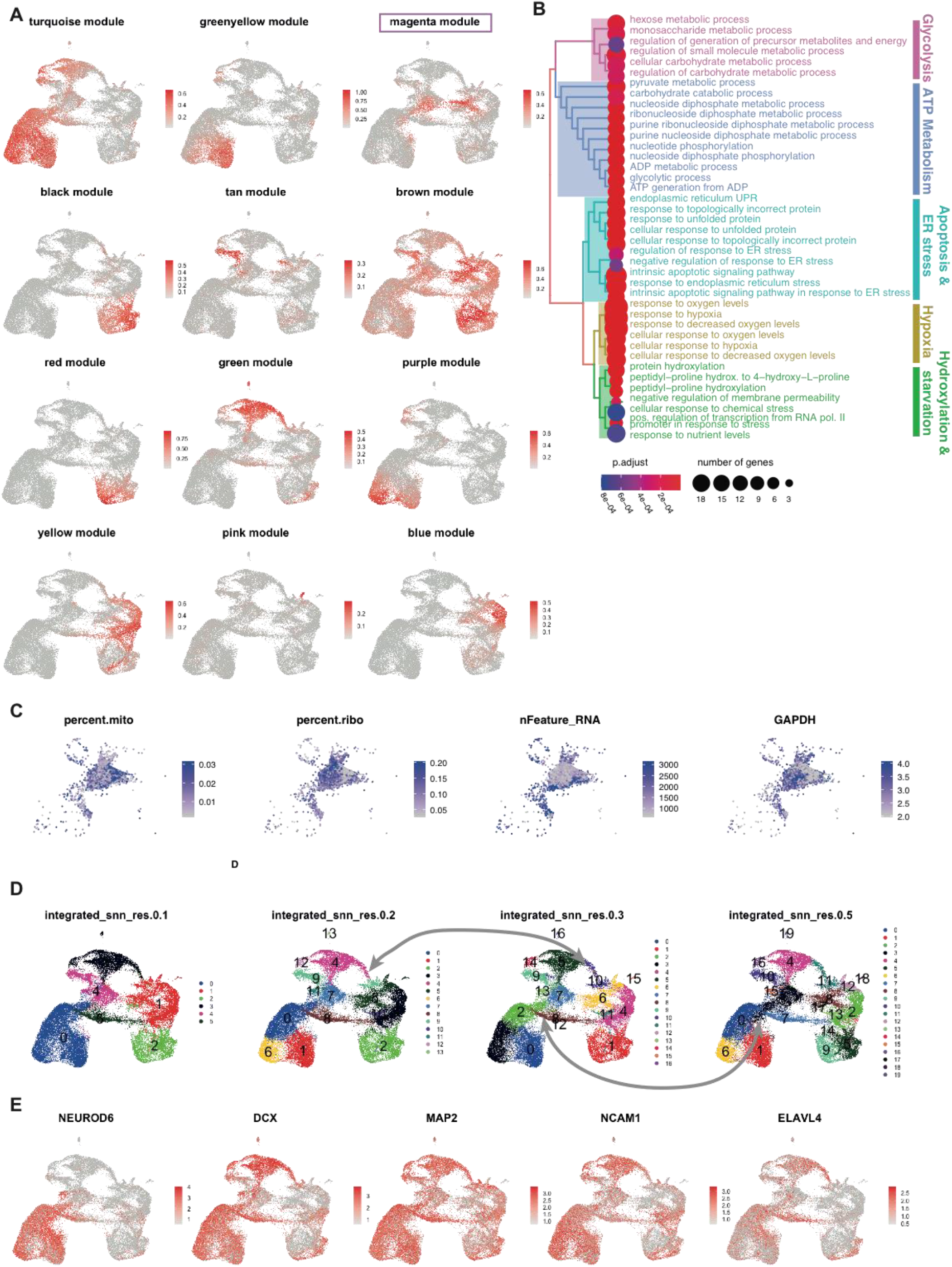
WGCNA modules the limitations of large cluster based analysis. (**A**) UMAP plots of 12 module-scores identified by single cell WGCNA. (**B**) GSEA of stress module (magenta), related to Fig 1G. (**C**) Heterogeneous composition of the ‘stressed-neuron’ cluster. The cluster shows salient and complimentary patterns of mitochondrial (I) or ribosomal (II) read fractions, as well as feature count (III). The expression of GAPDH (IV) signals the population of cells with high glycolytic scores. (**D**) Clustering resolutions 0.1 to 0.5 on integrated organoid dataset. Small resolution with large clusters merges stressed and unstressed cell types. With increasing resolution cluster boundaries change and make clear identification of stressed cell clusters difficult. (**D**) Expression of classic or pan-neural markers is diminished in stressed neurons. (**E**) Expression of classic or pan-neural markers is diminished in stressed neurons.

**Fig S2.**
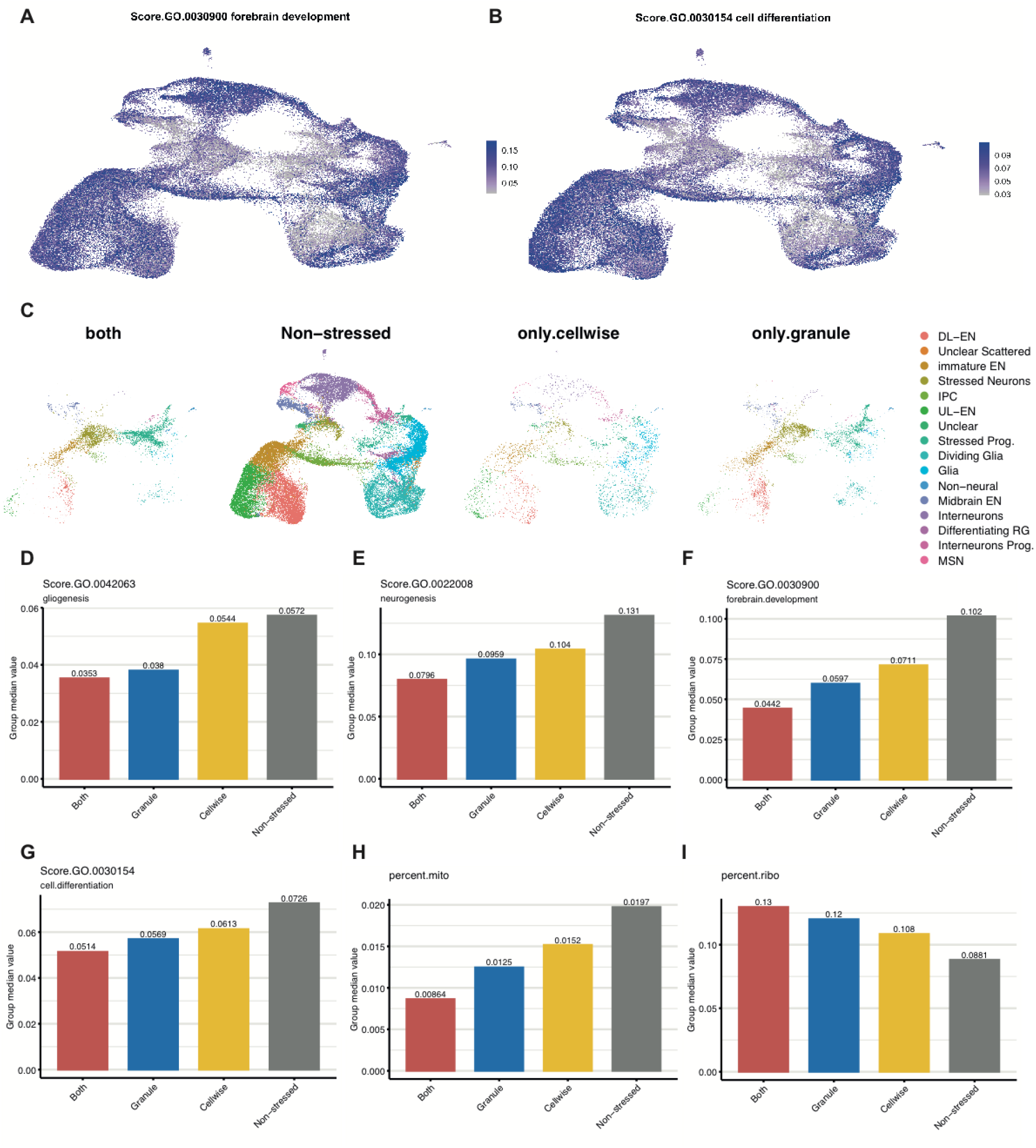
Differentiation related GO-terms are depleted in stressed cells\. Stressed clusters showed remarkably low pathway scores for **(A)** ‘forebrain development’ (GO:0030900) and for **(B)** ‘cell differentiation’ (GO:0030154). (**C**) UMAP comparison of stressed cells identified by Gruffi’s granular method (gSC) and stressed cells identified on single-cell scores (scSC). Cells are colored by clusters as in (**Fig. 1**), and separated by stress identification classes (identified by either, both or neither of the approaches). Class median values for gliogenesis (‘GO:0042063’, **D**); neurogenesis (‘GO:0022008’, **E**); forebrain.development (‘GO:0030900’, **F**); cell.differentiation (‘GO:0030154’, **H**); mitochondrial- and ribosomal mRNA content (**H, I**).

**Fig S3.**
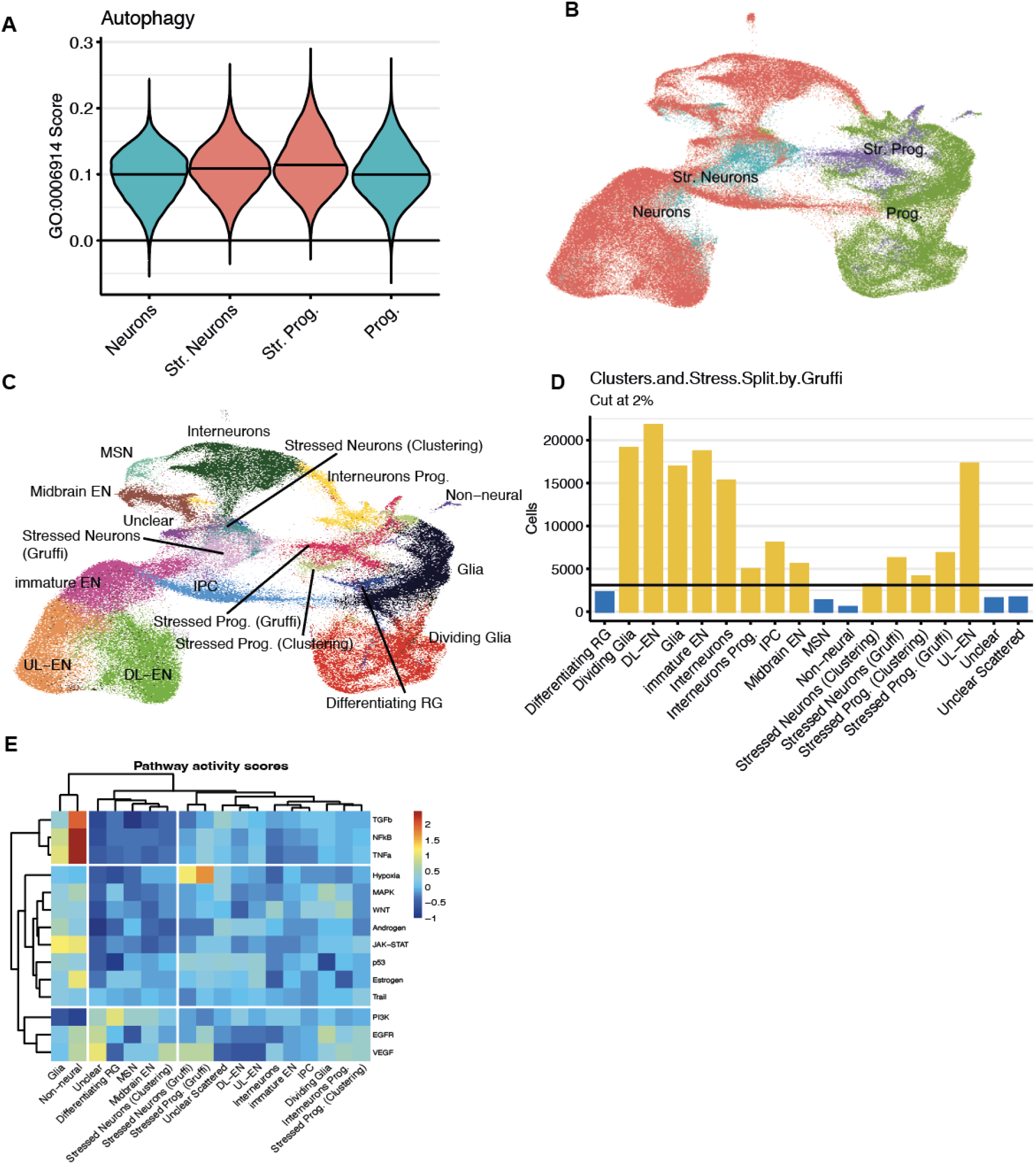
Mitochondrial and ribosomal balance and degradation pathways in stressed cells. (**A**) Fraction of mitochondrial reads (lect) per cell shows relative depletion in stressed neurons but not glia (right). Fraction of ribosomal reads per cell (middle) shows relative enrichment in stressed glia but not normal progenitors (right). (**B**) Fraction of mitochondrial reads is significantly higher in stressed cells (Kruskal-Wallis Test) (**C**) Fraction of ribosomal reads is significantly lower in stressed cells (Kruskal-Wallis Test) (**D**) Cluster median fraction of mitochondrial and ribosomal reads. (**E**) and (**F**) GO-terms score autophagy of mitochondrion (GO:0000422, (**E**)), but not autophagy (GO:0006914, (**F**)) is characteristic of stressed cells. (**G**) Cluster median expression of autophagy of mitochondrion (GO:0000422) and autophagy (GO:0006914). (**H**) GO-terms score ubiquitin-dependent ERAD pathways (GO:0030433) is specifically upregulated in stressed cells. (**I**) Cluster median expression of ubiquitin-dependent ERAD pathways (GO:0030433)

## Supplementary Table Descriptions

Table S1

A small, clearly distinct population of ribo-high cells was found to come from a single dataset, and constituted mis-patterned, non-neural tissue. This table quantifies the distribution of these ribo-high cells (>30% ribosomal reads) across datasets and clusters.

Table S2

Comparison of ribosomal, mitochondrial reads and stress scores in stressed and unstressed clusters.

Table S3

We calculated modules of co-regulated genes on the fetal reference dataset (Polioudakis et al., 2019), that were subsequently aggregated to correlate fetal and organoid data.

Table S4

Samarasinghe DGEA results and stressed-cell enriched coding genes. STRING DB’s GO - Biological Process enrichments on the above gene lists. Terms highlighted in Fig EV3C and permalink.

Table S5

We calculated differentially expressed genes enriched in excitatory neurons (first tab) and progenitors (second tab) in the fetal reference dataset (Bhaduri et al. 2020). The top 30 genes were selected for progenitor and excitatory neuron scores, respectively (third tab).

Table S6

Gene modules for plotting upper and deep layer neuron scores on pseudotime maturation. Gene modules were calculated with monocle3 and respective modules were selected.

## Methods

### Experiments

#### Stem cell culture

We obtained the “HPSI0114i-rozh_5“ (female) line from the HipSci catalog (Streeter et al. 2017), hiPSC cells were cultured following the [HipSci guidelines](https://www.hipsci.org/instructions/feeder-free-culture-protocol.pdf). We also grew organoids from the feeder-free human ES cell line (H9; WA09 from WiCell, Female). The two iPSC Lines SCCF – 177 (177J clone#8, female) and SCCF – 178 (178J clone#5, male) were generated at the IMBA Stem Cell Core Facility and are part of the IPSC Biobank. The study was approved by the local ethics committee of the Medical University of Vienna (MUV). After informed consent, a skin biopsy was taken from three healthy donors and fibroblasts were isolated for iPSC reprogramming. iPSC lines were generated using the Sendai virus (CytoTuneTM-iPS 2.0 Sendai Reprogramming Kit, Thermo Fisher Scientific) carrying the Yamanaka reprogramming factors OCT3/4, SOX2, c-MYC and KLF4 factors. All cell lines were used within 10 passages from last STR profiling and tested regularly for mycoplasma contamination. We additionally used the above cell lines (177 and 178) for the media comparison experiments (Fig. 4). The cell lines were evaluated and cultured as the HipSci lines. Briefly, cells were seeded on vitronectin (Stemcell Technologies, cat#07180) coated plates and fed every day with E8 essential media. Cells were passaged as single cells using Accutase (Sigma) with Revitacell cell supplement (1/100, Invitrogen, cat#A2644501), and grew until 70% confluency, then we replated. Cultured cell lines were routinely tested for mycoplasma contamination by PCR (Janetzko et al. 2014).

#### Organoid culture

Organoids were generated as described in (Esk et al. 2020). Briefly, 150 uL/well of Essential 8 media supplemented with RevitaCell (1/100) containing the corresponding cell suspension for 9000 cells/well were plated for each cell line using low attachment 96-well plates (Sigma CLS7007). Briefly, the protocol entailed the following steps: On day 3, media was replaced to Essential 8 media and from day 6 on, embryoid bodies were transferred to neural induction media (NI) and 200uL/well was exchanged every day. On day 10, when embryoid bodies are about 500-600um in thickness and neuroepithelium is evident, the aggregates were transferred to 10 cm dishes and embedded in matrigel (MG) droplets. On day 13 and 14, the media was changed to Improved Differentiation Media without ascorbic acid (Imp-A) containing 3 uM CHIR. After that, the media was replaced every 3-4 days. On day 19, the dishes were transferred to an orbital shaker. On day 25, the organoids were fed with Improved Differentiation media with ascorbic acid (Imp+A) and the media was replaced every 3-4 days. On day 40, the two different culture methods (Brainphys & Imp+A) diverged. The “Imp+A” organoids were further cultured in Imp+A supplemented with 1%MG, BDNF (20ng/mL), GDNF (20ng/mL) db-cAMP (1mM). The “Brainphys” (BP) organoids were gradually transitioned to BP media in 3 feeding steps: 75%-25%: 50%-50%, 25%-75% (Imp+A & BP). From that point they were cultured in BP supplemented with 1%MG, BDNF (20ng/mL), GDNF (20ng/mL) db-cAMP (1mM).

#### Media composition

##### NI media

Neural Induction medium consisting of DMEM/F12 (Thermo Fisher Scientific) with 1% N2 Supplement (Thermo Fisher Scientific), 1% MEM-NEAA (Sigma Aldrich), 1% Glutamax (Thermo Fisher Scientific) and 1 ug/ml Heparin. **Imp-A**: of 50% DMEM/F12 (Thermo Fisher Scientific), 50% Neurobasal (Thermo Fisher Scientific), 0.5 % N2 Supplement (Thermo Fisher Scientific), 2% B27 - Vitamin A (Thermo Fisher Scientific), 1% Glutamax (Thermo Fisher Scientific), 0.5% MEM-NEAA (Sigma Aldrich), 50uM 2-ME solution, 1 % Penicillin/Streptomycin (Sigma Aldrich) and 0.025% Insulin solution (Sigma Aldrich). **Imp+A (HN)**: Imp-A with 2.5 mM Ascorbic Acid, 2g/l Bicarbonate (Sigma Aldrich). **BP (LN)**: BrainPhys Neuronal Medium (Stem Cell Technologies), 2% B27+A (50X, Thermo Fisher Scientific), 1% N2 supplement (Thermo Fisher Scientific), 200 nM Ascorbic Acid (Sigma Aldrich), 0.2% CD Lipid Concentrate (Thermo Fisher Scientific), 7.4% glucose, and 1% Penicillin/Streptomycin

#### Single-cell sequencing

Organoids were cultured to 120 days, then washed in PBS and dissociated using the gentleMACS Dissociator (Miltenyi Biotec) in program NTDK1 using the enzyme mix: Trypsin (Sigma Aldrich)/Accutase (Sigma Aldrich) (1:1) containing 10 U/ml DNaseI (Thermo Fisher Scientific). The washed cell suspension was passed through a 70μm cell strainer.

In the newly generated datasets, we pooled cells from samples from 4 different genotypes and were combined in a lane (other cell lines used for other purposes). We additionally used sample barcoding using lipid-anchor barcoding following instructions as in (McGinnis et al. 2019) with reagents kindly provided by the authors, but we relied on SNP-based cell line demultiplexing as described in (Kang et al. 2018) (described in the following section) and sample barcoding was not used.

Cells were counted and the suspension was loaded onto a Chromium Single Cell 3′ B Chip (10x Genomics, PN-1000073) and processed through the Chromium controller to generate single-cell GEMs (Gel Beads in Emulsion). scRNA-seq libraries were prepared with the Chromium Single Cell 3′ Library & Gel Bead Kit v.3 (10x Genomics, PN-1000075).Ready 10X libraries were sequenced paired end (R1:28, R2: 89 cycles) on NovaSeq (Illumina).

### Data Analysis

#### Public Datasets

We used the following public datasets for this study: dbGaP Study Accession: phs001836.v1.p1. (Polioudakis et al. 2019; de la Torre-Ubieta et al. 2018); ENA PRJEB33917 (Kanton et al. 2019); GEO GSE132672 (Bhaduri et al. 2020); EGA EGAD00001006332 (Eichmüller et al. 2022); GEO GSE129519 (Velasco et al. 2019).

#### Cell line demultiplexing

For pooled 10x GEX libraries the donor cell-line of the assayed cells was determined by genotype-based demultiplexing using souporcell (Heaton et al. 2020). The pipeline was run with default settings, providing all donor genotypes through the known_genotypes parameter, and providing the cellranger bam, the cellranger filtered barcodes file, and the reference fasta as input. Donor genotype vcfs were pre-generated using HaplotypeCaller from the Genome Analysis Toolkit (GATK) v4.1.2.0 on bwa mem 0.7.17 aligned WGS reads following the nf-core/sarek v2.5.1 pipeline. WGS reads were obtained from respective sources: (a) SRA database for H9/SRR6377128, (b) from the ENA database for the HIPSCI line rozh_5/ERR1871976 or (c) WGS data generated by the IMBA stem-cell facility for SCCF – 177 (177J clone#8, female) and SCCF – 178 (178J clone#5, male).

#### Single-cell analysis

We first aligned reads to the GRCh38 human reference genome with Cell Ranger 3.1 (10x Genomics) using pre-mRNA gene models and default parameters to produce the cell-by-gene, Unique Molecular Identifier (UMI) count matrix. UMI counts were then analyzed in R, using the Seurat v4. We filtered for high quality cells based on the number of genes detected (>500). Thereafter, expression matrices of high quality cells were normalized (“LogNormalize”) and scaled to a total expression of 10K UMI for each cell. Regression of variables at this step did not improve clustering results, hence no variables were regressed nor removed.

##### Non-neuronal cell exclusion

Before integration datasets were checked for quality, as certain IPS lines are prone to misdifferentiation. As a consequence, multiple datasets included non-neuronal cells that would interfere with the CCA integration, henceforth we applied initial filtering for CNS cells. To exclude non-neuronal cells we used a recently published fetal organ atlas (Cao et al. 2019). Processed data was downloaded and cell type annotation was modified to reflect major cell types for a basic classification. All CNS cell types were grouped together under one annotation to determine properly specified clusters. Next, an xgboost classifier was trained to distinguish major cell types on the RNA assay data using the top variable genes of the fetal dataset with parameters determined by cross-validation. This classifier was applied to each dataset: 1. Datasets were pre-processed individually and clustered in UMAP space; 2. The expression of the RNA assay was used to classify each cell according to the cell groups of the training dataset; 3. Classification was summarized per cluster and all clusters that were not classified as CNS cells were filtered out. The cleaned datasets were used for CCA integration.

To establish the maximal mitochondrial-, and ribosomal RNA fractions, we plotted these against feature counts and each other, and set thresholds to remove extreme outliers. A group of cells showed a distinctly high ribosomal fraction (>30%). We found that these cells correspond to one cluster coming from one dataset (Velasco organoid 21, Table S1) and are non neural in gene expression. We used the same threshold value for maximal mitochondrial read fraction for simplicity.

##### Downstream Analysis

Variable genes were identified by Seurat’s FindVariableFeatures implementation (“FastLogVMR”). Next, we aligned and merged sequencing libraries by Seurat’s canonical correlation analysis or CCA (dimensions: 50) (Butler et al. 2018) using the intersection variable genes across datasets.

Next, principal components were calculated on the variable genes, and the first 50 components were then used to calculate UMAP coordinates. For clustering we used Seurat’s implementation of snn/Louvain clustering. Therein, we first calculate the k-nearest-neighbor (knn) graph of cells in PCA-space (dimensions:50). Based on Jaccard similarity scores on the knn graph the shared nearest neighbor (snn) graph is computed. Louvain clustering on the snn graph identified clusters of cells. Differentially expressed genes were identified by Wilcoxon-test, and filtered for p-values below 0.001, and fold change larger than 2.

We found a group of 1304 interneurons that formed a separate cluster on the very top of the UMAP (Fig 1). These constituted 7.61% of all interneurons and were 94% originating from the Kanton S3 dataset. Both interneuron clusters showed similar expression of classic interneuron markers, and pairwise differential gene expression analysis showed no meaningful differences. Therefore, we lumped these 2 clusters together

#### Integration of organoid and fetal data

We obtained raw data for fetal cortical single cell datasets covering age comparable to organoid datasets (de la Torre-Ubieta et al. 2018; Polioudakis et al. 2019), pre-processed and analyzed it, the same way as we did for the organoids. The individual sequencing lanes were merged per fetal datasets and integrated with Seurat, as before. The integrated organoid dataset was uniformly downsampled to 24211 cells, to match the total sample size of the fetal datasets. The resulting individual (original) datasets were then reference-integrated to the fetal dataset as follows: First, 3000 integration anchors were computed with Seurat’s SelectIntegrationFeatures() and FindIntegrationAnchors() functions, where the fetal datasets were defined as reference. By default the integration by IntegrateData() was performed using CCA, setting parameter k.weight to 50. Further steps, such as the determination of variable features, scaling, the computation of PCA and UMAP embeddings and the SNN Graph were performed as for the organoid integration.

#### Individual analysis of Velasco 7 dataset (Fig EV2)

The dataset was filtered for high quality cells with a higher gene count than 1000 and analyzed using Seurat as described above including log-normalization, scaling, the computation of the 2000 most variable genes, PCA and UMAP embedding computation.

### The Gruffi package

The Gruffi package contains all functions for the identification, inspection and filtering of stressed cells using command line or graphical user interface (Shiny app). Gruffi functions encompass the following major steps as in (Fig.2B): (1-3) Accession of GO-term gene sets and single-cell stress scoring; (I-III) Data partition into granules and small-granule reclassification; (4) Aggregate score calculation per ensemble; (5) Automatic estimation of stress threshold, with possible manual adjustment and inspection; (6) Stressed cell assignment and filtering.

#### Single-cell scoring

We defined specific GO-terms relevant for functional processes in stress and differentiation. Gene lists for each GO-term were downloaded from Ensembl via BiomaRt (Durinck et al. 2009) and intersected with detected genes. Alternative database access is also implemented (see R-package documentation). We then generalized a widely used cell-cycle scoring method based on aggregated gene set activity (Tirosh et al. 2016), and used its implementation in Seurat (*AddModuleScore*). Briefly, in the *AddModuleScore* function the following steps have been implemented as in the original: (A) Take a target set of genes; (B) Calculate their average expression; (C) Create control sets of genes. The control gene-sets are used to control for the cell-to-cell variability in quality and depth. To create the control sets, first all genes are binned by expression levels (25 bins), then for each gene in the target set, randomly select 100 genes from its expression bin, and finally (D) subtract the average of control from each target gene. The expression binning is important, because expression levels affect the variability of gene expression (Tirosh et al. 2016). Gene lists of ‘glycolytic process’ (GO:0006096) and ‘response to endoplasmic reticulum stress’ (GO:0034976) were downloaded and intersected with detected genes, then used to evaluate stress state.

#### Data partitioning and reclassification

Gene detection in single cells is noisy. To overcome this noise, we grouped cells into small aggregates by high-resolution snn-clustering (as in the manuscript, using the algorithm of Seurat). Gruffi’s *aut*.*res*.*clustering()*, finds an optimal granule resolution with a median of 100-200 cells per granule (cluster). Next, clustering is performed at the determined resolution, resulting in cells separated into 100’s of granules, depending on the size of the dataset. Finally, all cells in granules with <30 cells are reassigned to the nearest cluster center in the 3D UMAP space (Euclidean distance) using *reassign*.*small*.*clusters()*.

#### Thresholding and stress annotation

Finally, the average GO-scores for each granule were calculated, and stress level per granule was evaluated. We propose two possible methods to estimate an upper threshold for the assignment of stressed granules for one GO term. (a) Determining manually, based on the expression of stress genes and stress score values on UMAP. Based on these, an empirical quantile (90% if observing 10% stressed cells) as threshold can be assigned. (b) Automatic stress threshold estimation by *Shiny*.*GO*.*thresh()*. For this, we refer to cell number normalized, mean GO scores per granule. In the following this will be referred to as granule score. We assume that: b1) The granules consist of a statistically sufficient number of cells. b2) GO scores of non-stressed cells independently follow the same unknown distribution. b3) GO scores of stressed cells are significantly higher than GO scores of non-stressed cells. b4) The dataset consists of more non-stressed than stressed cells. Assumptions b1) and b2) together with cell number normalization allow us to use the central limit theorem and hence fit a normal distribution to the granule scores of non-stressed granules. Based on b3) and b4) we conclude that respective non-stressed GO-scores are small, and the mean of the normal distribution can be estimated by the median of granule scores. The standard variation of the normal distribution is now estimated only w.r.t. to GO scores smaller than the median of granule scores. Now the theoretical 99% quantile of the fitted normal distribution can be computed and used as an upper threshold.

Assumption b1) is fulfilled by the automatic clustering resolution search and the reassignment of cell granules with less than 30 cells. Although we based our analysis on thresholds retrieved by this method, since we cannot assure that assumptions b2) to b4) hold true, we highly recommend further inspections and refinements of suggested thresholds in any case. To do so, we propose to visually monitor further manual adjustments via the implemented Shiny App interface.

When considering a combination of GO terms, e.g., response to Endoplasmic Reticulum Stress and Glycolytic Process, we combine the respective thresholds such that a cell is assigned as stressed if either upper threshold is crossed. In case one additionally wants to include a GO term for non-stressed cells, e.g., gliogenesis, the above threshold method can be applied, too, but in this case granules with a score higher than the threshold are assigned as non-stressed. Finally, based on the thresholds on each score, stressed cells are annotated and can be excluded from the dataset.

### Other analyses

#### Protein-protein interaction maps

We selected all genes enriched in either stress clusters and jointly ranked them by descending log2FC. We selected the top 150 coding genes, and visualized the ‘high-confidence’ connected component of the protein protein interaction network using the STRING database (v11.5) (Szklarczyk et al. 2019), links denoting the confidence of connection (permalink: bxso1NJafq8R).

#### Pathway visualization using ShinyGO and KEGG

The top 150 coding genes (as above) were provided for ShinyGO v0.741 ((Ge, Jung, and Yao 2020) with default parameters and the background gene list of all 26439 detected genes (from the RNA assay). ShinyGO’s visualized enriched KEGG pathways using Pathview and relevant pathways were selected.

#### Comparison of granular and single-cell scoring

For granular scoring, we used the annotation and approach from (Fig 2). For single-cell scoring, we used the exact same approach, but skipped the granule average calculation of stress scores, and instead we calculated the stress threshold on single-cell scores. For fair comparison, we adjusted the quantile cutoff parameter in the single-cell scoring so that it results in a similar number of stressed cells, as in the granular approach. We then took the symmetric difference of these to find cells only flagged by either, but not both methods. Group median values for were plotted for the four categories (Both, gSC, scSC, Non-stressed)

#### The separation of stressed neurons

Stressed cells clearly separated into two major groups, as also seen in **(Fig 1)**. Therefore, we separated Gruffi’s classification into two categories. Low resolution clustering (res.0.1.ordered) separated glia (cl.1-2) from neurons (remaining clusters) both better (less clustering artifacts) and simpler than higher resolutions (0.3, 0.5). Intersecting this binary annotation with Gruffi’s stress annotation (T, F), separated cells into the four clusters: Neurons, Stressed Neurons, Stressed Progenitors, Progenitors, visible in (**Fig S3B**).

#### Progeny pathway activity scoring

Progeny pathways scoring was performed as in vignette, with the following parameters top 200 genes. To visualize the differences between stressed cells identified by typical clustering (Fig. 1B) and Gruffi (Fig. 2F), we separated “Stressed Neurons” and “Stressed Prog.” clusters into subsets identified, or not identified by Gruffi, yielding four groups: “Stressed Prog. (Clustering)”, “Stressed Prog. (Gruffi)”, “Stressed Neurons (Clustering)”, “Stressed Neurons (Gruffi)” **(Fig S3C)**. As progeny failed to run on the full object, we randomly downsampled the full dataset to 33.3% of the cells (>50K cells). We visualized the scores using pheatmap with ward.D2 hierarchical clustering and separated the 3 most distinct clusters. For (**Fig 3B**) we displayed all clusters >2% of all cells **(Fig S3D)**, and all clusters are displayed in **(Fig S3E)**.

#### Choroid Plexus scoring and stress identification in Samarasinghe et al

We obtained the Seurat R object of (Pellegrini et al. 2020) from cells.ucsc.edu and performed DGEA by Wilcoxon test in Seurat. We used the clustering presented in the paper, and contrasted “mature choroid plexus” to all other clusters. We calculated a choroid plexus score from the resulting 192 genes (log2fc>1, p.adj<0.01, pct.expr>33%, **Supplementary Table S5**) and provided this to Gruffi as a negative score (like gliogenesis). We then calculated differential gene expression on stressed cells vs. non-stressed cells, as identified by Gruffi. The resulting 16 genes (log2fc>1, p.adj<0.01) were then analyzed in STRINGdb as before (permalink: bwEKXY7CP0p8).

#### Neural and glial identity scores for cell type specification

As previously (Bhaduri et al. 2020), we grouped all neural or progenitor classes to define the respective tow signatures in fetal samples, based on DGE analysis. After intersection with genes that are detected in organoid datasets, the top 30 genes were used for EN, or progenitor signatures (**Supplementary Table S6**). From these, per-cell subtype scores were calculated using the AddModuleScore() function of Seurat. XY-scatter density plots were drawn by plotting a progenitor score and a neuronal score on the X and Y axis respectively. The cells were colored based on cell type reflecting progenitors, neuronal cells and intermediate progenitors (IPCs), that would physiologically be an intermediate state.

#### Pseudotime analysis of maturation

For analysis of the ‘dorsal lineage maturation’ datasets of pairwise H- and L-medium organoids were integrated as outlined above. The datasets were then transferred to monocle3 and UMAP was calculated with three dimensions. The trajectory graph was constructed on the three dimensional dataset from progenitors to mature neurons. In order to compare the maturation of gene expression modules of co-regulated genes were calculated with the *find_gene_modules()* function. A module for Upper Layer and Deep Layer Neurons was selected for each mature dataset (**Supplementary Table S7**). To plot maturation in the different datasets each cell was plotted along pseudotime (x) versus the expression of the respective score (y).

## Footnotes

1 ALT: No sign of general neural maturation problem and misspecification in organoids

